# Hallucination of closed repeat proteins containing central pockets

**DOI:** 10.1101/2022.09.01.506251

**Authors:** Linna An, Derrick R Hicks, Dmitri Zorine, Justas Dauparas, Basile I. M. Wicky, Lukas F. Milles, Alexis Courbet, Asim K. Bera, Hannah Nguyen, Alex Kang, Lauren Carter, David Baker

## Abstract

In pseudocyclic proteins such as TIM barrels, β barrels, and some helical transmembrane channels, a single subunit is repeated in a cyclic pattern, giving rise to a central cavity which can serve as a pocket for ligand binding or enzymatic activity. Inspired by these proteins, we devised a deep learning-based approach to broadly exploring the space of closed repeat proteins starting from only a specification of the repeat number and length. Biophysical data for 38 structurally diverse pseudocyclic designs produced in *E. coli* are consistent with the design models, and two crystal structures we were able to obtain are very close to the designed structures. Docking studies suggest the diversity of folds and central pockets provide effective starting points for designing small molecule binders or enzymes.

## Introduction

Native cyclic repeat proteins carry out a very broad array of functions in biology. For example, the triosephosphate isomerase (TIM) barrel^1^, which consists of eight **ɑ**/β repeats that close to form an 8-stranded β barrel surrounded by an outer ring of helices, is the most prevalent protein fold for enzymes. Single-chain cyclic structures formed by repeating units have considerable advantages: at the center is a pocket into which side chains from each repeat unit extend, and because it is a single chain, the sequence lining (and local structure) can be fully asymmetric. De novo protein design has been used to create repeat proteins which do not close^2,3^, and closed TIM barrels^4^, parametric bundles^5^, and all ɑ helical toroids^6^. However, a general method to broadly sample repeating cyclic structures without specifying the overall architecture or lengths and positions of the secondary structures has thus far been missing.

We sought to develop a general approach for overcoming these limitations to enable generation of a wide range of new cyclic protein scaffolds with central cavities for small molecule binder and enzyme design. We reasoned that recently developed deep network-based protein hallucination methods^7^, which optimize sequences for folding to specific structures, without requiring specification of what that structure is, could be extended to broadly sample cyclic repeat protein structure space given only the repeat unit length and number of repeats.

## Results

### Hallucination and Sequence Design of Pseudocycles

We developed a sequence space Markov Chain Monte Carlo (MCMC) optimization protocol (**Figure 1a**) which, given specification of the length *L* and number *N* of repeating units, first generates a random amino acid sequence of length *L* and tandemly repeats it *N* times. We sampled *N* from 2 to 7, and *L* from 15 to 78, with a max protein length of 156 amino acids. The protocol then optimizes this sequence by making 1-3 random amino acid substitutions at a random position in one repeat unit, propagating these mutations to all repeat units, evaluating the extent to which it encodes a cyclic repeating protein structure, and finally accepting or rejecting the substitutions according to the standard Metropolis criterion. To evaluate sequence folding to a cyclic structure, we used AlphaFold2^8^ (AF2) with a single sequence as an input and 3 recycles to predict the structure, and subsequently evaluated the extent of closure by extrapolating helical parameters from the rigid body transforms between successive repeat units: closed structures are those with near zero rise along the helical axis, and rotation of 360/*N* degrees about the helical axis. To guide the MCMC trajectories, we supplemented the closure score with AF2 confidence prediction metrics (see Online Methods Protein generation and sequence design pipeline). We found that after only a few hundred steps (see **supplementary Figure. S1**), most MCMC trajectories optimizing for this combined score converge on sequences which are predicted to fold with high confidence into closed cyclic structures (**Figure. 1b**).

**Figure 1.**
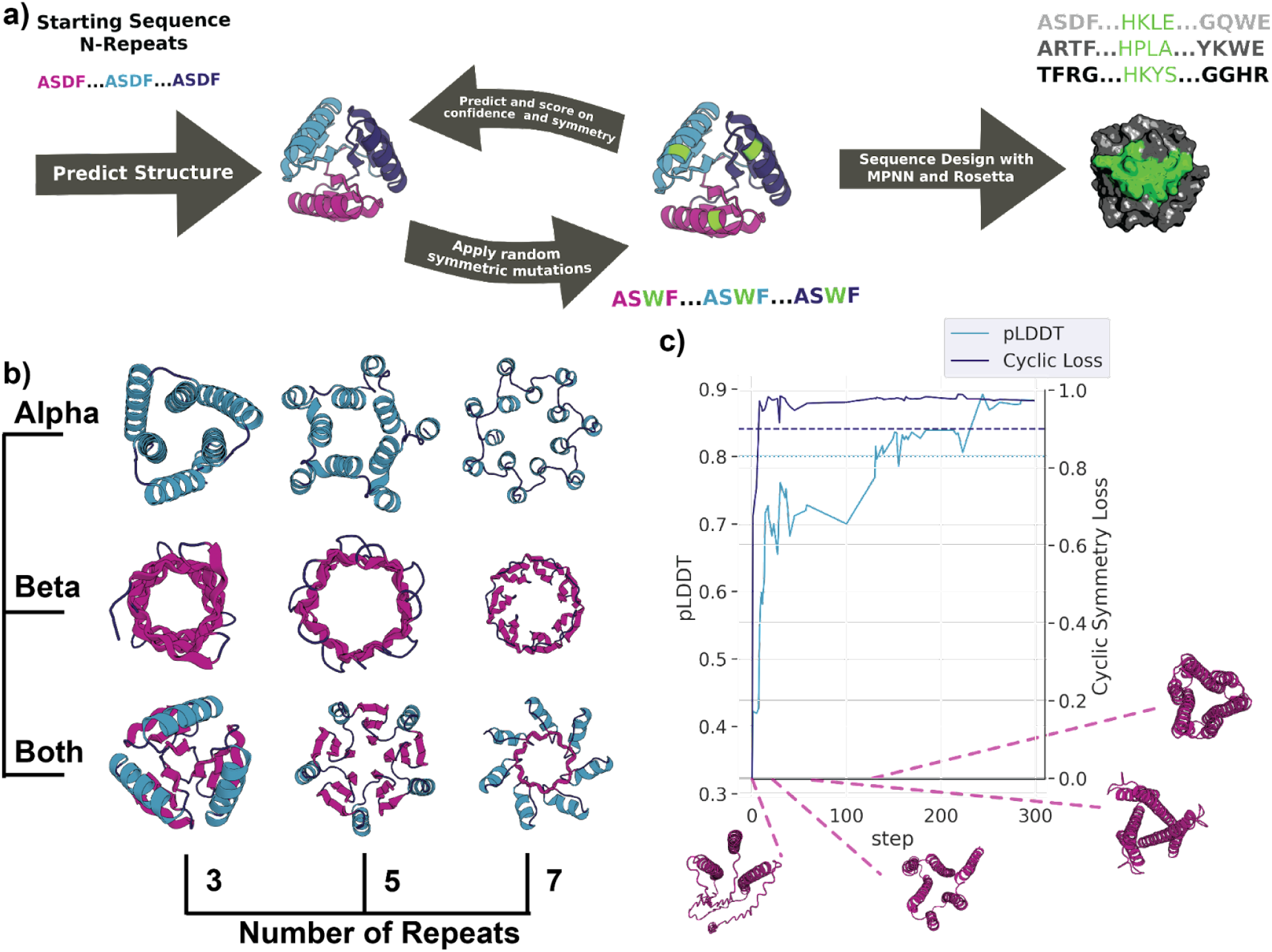
Pseudocyclic protein design. a) Schematic representation of the scaffold hallucination and design pipeline. b) Selected output proteins featuring 3, 5 or 7 repeats, and all-**ɑ** (Alpha), all-β (Beta) or mixed **ɑ**/β (Both) topologies. c) Representative design trajectory showing the optimization of pLDDT (teal) and cyclic loss (dark blue) over 300 steps, with dashed lines indicating our selected score cutoffs. Protein structure cartoons are snapshots at indicated steps in the trajectory; loop, sheet, and helix regions are colored in dark blue, magenta, and teal, respectively.

While this cyclic hallucination procedure generated a very wide array of new cyclic backbones, the actual amino acid sequences contained sub-optimal features such as large hydrophobic surface patches and poor secondary structure sequence agreement (see **supplementary Figure. S2 and S3**). A limitation of the hallucination procedure, as with any activation maximization procedure which optimizes over the inputs to a neural network, is generation of adversarial examples by overfitting. Hallucination studies on cyclic oligomer design showed that while AF2 generated sequences were rarely soluble, redesign of the hallucinated backbones with ProteinMPNN yielded soluble proteins with desired structures (basile/lukas 2022 unpublished). Hence, we used ProteinMPNN (Justas 2022 unpublished) to design new sequences given the hallucinated backbones (see Online Methods Protein generation and sequence design pipeline) without requiring sequence repeat symmetry, which resulted in sequence-asymmetric final designs (**Figure. 1c**). Finally, we used RoseTTAfold^9^ (RF) and AF2 to evaluate the extent to which the designed sequences encoded the designed structures (**supplementary Figure. S4**).

We obtained a total of 21,021 designs strongly predicted to fold to the intended structures. We refer to these designs as ‘pseudocycles’ because their backbones have near cyclic symmetry (except for the break between the C and N termini) but the sequence is asymmetric. The 21,021 designed pseudocycles span a very wide range of topologies containing all **ɑ**, **ɑ**/β, and all β subdomains (see **Figure. 1b** and **supplementary Figure. S4**). In some of the designs, the repeat units form compact domains which interact with neighboring units through relatively small interfaces, while in others the repeat units are more intertwined (**Figure. 3 a & b** and **supplementary Figure. S5**).

### Experimental Characterization of Selected Pseudocycles

We selected 96 designs with a range of repeat unit numbers, lengths, and secondary structure compositions for experimental characterization, focusing on designs containing designable pockets and structural folds rarely found in the Protein Data Bank (PDB) (**Figure 2**). These proteins have sequences and structures quite different from those in the PDB, with median BLAST e-value of 0.018 and TMscore’s between 0.33 - 0.87 and average of 0.54 (see **supplementary Figure. S6** and **supplementary Table. S1, S2**). Following expression in *Escherichia coli*, we found that 81 of the 96 designs were soluble and 38 of these 81 soluble designs were well-expressed and had circular dichroism (CD) spectra which indicated that the proteins were well-folded with overall secondary structure content consistent with the design models (**Figure. 2** and **supplementary Figure. S7**). 17 of these 38 designs were monomeric and monodisperse, while another 15 were polydisperse with a majority monomeric population. (**Figure. 2** and **supplementary Figure. S8**).

**Figure 2.**
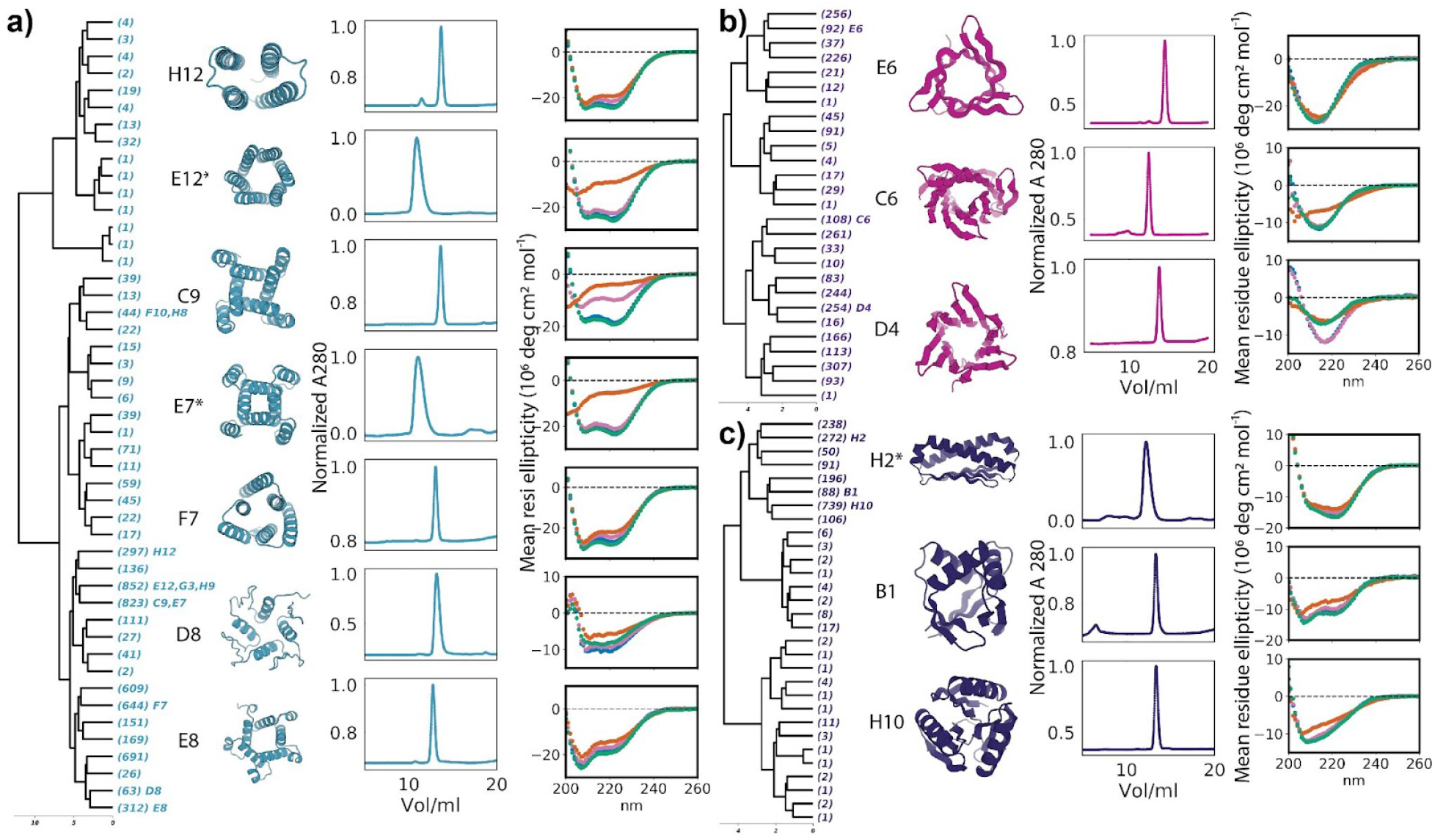
Biophysical characterization. Left panel in a, b, c: hierarchical clustering of designed pseudocycles. The number of sub-branches are indicated in brackets. 2nd panel: cartoon diagrams of designs selected for experimental characterization; identifiers indicate position in dendrograms. Third panel: SEC trace. Protocols are described in the supplementary material Expression and purification of selected proteins. Proteins prepared following protocol1 are marked with star(*). Fourth panel: CD spectra at different temperatures (25 ° in blue, 55 ° in orange, 95 ° in pink, followed by refolding at 25 ° in green). a) **ɑ** helical topologies (colored teal), b) β sheet topologies (colored magenta), c) mixed **ɑ**/β topologies (colored dark blue).

We were able to solve crystal structures of two designed pseudocycles. In both cases, the crystal structures show closed repeat structures very similar to the computational design models (**Figure. 3** and **supplementary Table. S3**). The first structure is a simple 4-helix bundle with pseudo-C2 symmetry formed by duplication of a helix hairpin repeat (**Figure. 3a**); the closest structure in the PDB (5OXF) has a TMscore of 0.65. The design model is very accurate with 0.8 Å CA-root-mean-square deviation (Ca-RMSD) to the solved structure. The second structure is a more complex pseudo-C3 symmetric protein with a repeated EEHE fold (**Figure. 3b-d**). The interface between repeat units contains seven buried hydrophobic residues contributed by a helix/strand motif from one repeat packing into a groove formed by three strands from the next repeat unit (**Figure. 3c**). According to the electron density map, a suite of water mediated hydrogen bonding networks was formed between the three center strands and water molecules occupying the central cavity (**Figure. 3d**). The design model was again very accurate, with 0.5 Å Ca-RMSD to the solved structure. There are no structures currently in the PDB with global similarity to this design (closest TMscore of 0.38 to 1U7Z). Both solved structures contain potential central pockets for functionalization (**Figure. 3a, b**).

**Figure 3.**
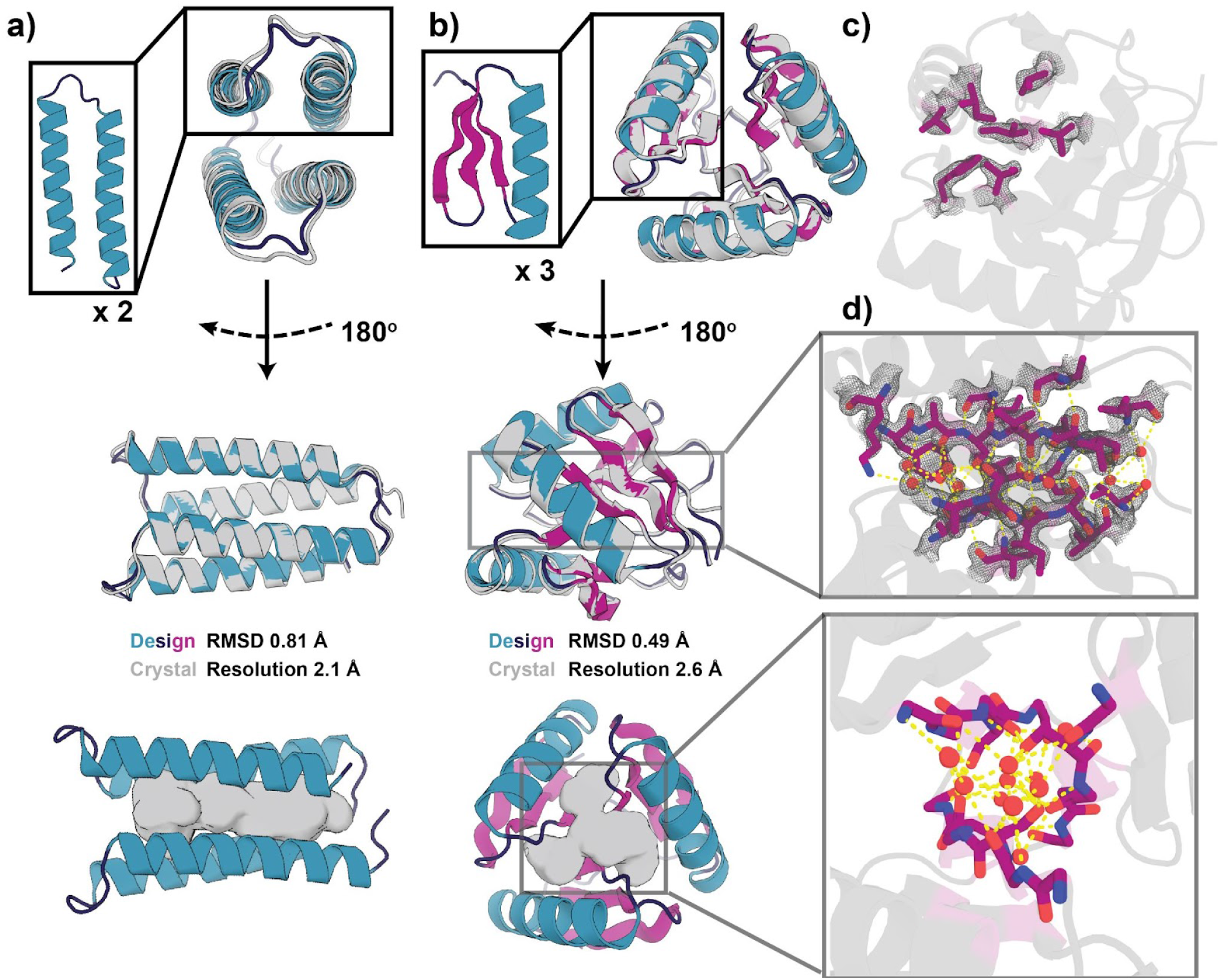
X-ray crystal structures of the designs are very close to the computational models. Crystal structures of the two-repeat design H12 (a) and the three-repeat design H10 (b-d) are shown in gray cartoons, the loop, sheets, and helix of the design are shown in dark blue, magenta, and teal, respectively. Central pockets in the designs are shown in gray (a, b). The secondary structure interface (c) and the center water mediated hydrogen bond network (d) of the refined crystal structure of design H10 are shown using sticks. The electron density map of the interface and the center hydrogen bond network and water are shown in gray mesh. The oxygen, nitrogen, and carbon are colored in red, blue, magenta in c and d; hydrogen bond networks are shown in yellow dashed lines, and water molecules in red spheres.

### Small Molecule Docking for Pseudocycles and other Scaffolds

To investigate the potential of the designs to harbor ligand binding pockets, we clustered 21,021 designed pseudocycles based on structural similarity (see supplementary information for clustering) into 9839 clusters. For each of the 9839 cluster centers, we carried out RIF docking^10^ and pocket design calculations with 19 ligands with diverse sizes, shapes, and chemical properties (see **supplementary Figure. S9**)^11^. For each of the 19 ligands, we also carried out RIF docking and design calculations for 2787 single-chain native small molecule-binding proteins from the PDB (PDBBind^12^) and 1000 previously published de novo designed NTF2-like proteins^13^ (see supplementary information for small molecule docking and binder design experiments). Docks of ligands to scaffolds were designed using Rosetta sequence design suite^14^ and Rosetta metrics were generated for evaluation of protein-small molecule binding interface quality^11^. The scaffolds most suitable for each ligand were picked out based on Rosetta metrics^11^. Examples of designed binding sites for several diverse ligands to their most-suitable pseudocyclic proteins are shown in **Figure. 4a**. We found that for most ligands, the most designable binding sites were obtained with the pseudocycle scaffolds (**Figure. 4b and c**, see supplementary information for small molecule docking, binder design, and binder evaluation experiments), likely because of the great variety of binding site shapes (see **supplementary Figure. S10**) and sizes, and CA-CB vector orientations surrounding the pocket, enable design of plausible binding sites for almost any ligand.

**Figure 4.**
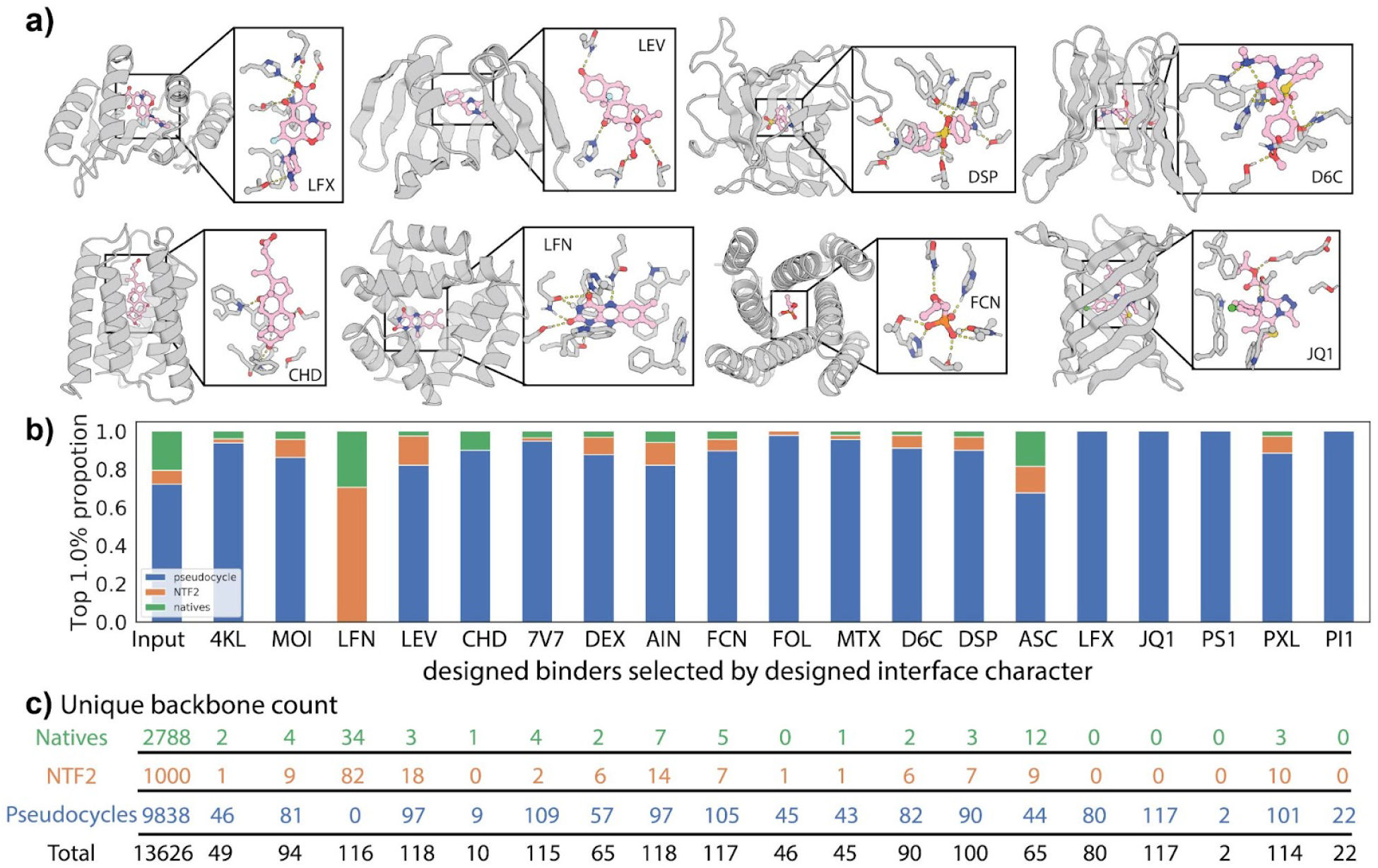
The central pockets can accommodate a wide range of small molecule ligands. a) Examples of computationally designed binding interactions for diverse ligands in pockets of diverse pseudocyclic scaffolds. Designed proteins are shown as gray cartoons/sticks and the ligands are shown in pink sticks. Oxygen, nitrogen, phosphorus, and chlorine elements are colored in red, blue, orange, and green, respectively. b) Barplots showing the composition of input scaffolds (pseudocyclic designs, natives from PDBBind database, and designed NTF2’s from Basanta et al^13^) and subsequent composition of best small molecule binders (each scaffold may generate as many as 30 binders) for diverse ligands. c) The numbers of unique backbone scaffolds selected based on the top 1% designed interface character from each type of scaffold are listed.

## Discussion

Our results further illustrate the power of deep network hallucination to explore the space of possible protein structures given only general specifications of structural features – in this case the number and length of the repeating units, and the constraint that the repeat units close on themselves to form an overall structure with cyclic symmetry. The approach generates a wide variety of structures strongly encoded by their amino acid sequences (as evaluated with RF and AF2, and further indicated by the close agreement between the crystal structures and design models), with between 2 and 7 repeat units (N), repeat lengths of 15 to 78 (L), and all **ɑ**, **ɑ**/β, and all β folds up to a max length of 156 amino acids. The sequences of the designs are unrelated to those of naturally occurring proteins, and while some structures resemble naturally occurring proteins, many have novel tertiary structures.

The de novo design of ligand binding proteins and enzymes is still in its infancy. Two approaches have previously been used. The first is to redesign naturally occurring proteins^15^, which is often limited by the relatively low stability of native proteins and the complexity of both the structure and sequence-structure relationships. Designs obtained with this approach are thus often unstable or have unpredictable structural changes. The second approach is to start from robust de novo designed scaffolds lacking features that are difficult to control, such as long loops, and have better understood sequence-structure relationships. This approach has been limited by the lack of diversity in available de novo designed scaffolds^10,13,16^ when compared to the diversity of structures in the PDB. The large set of pseudocycles with central binding pockets described here combines the diversity of native protein scaffolds with the stability and robustness of de novo designed proteins, and our RIF docking and Rosetta design calculations suggest that the designed pseudocycles provide better starting scaffolds for small molecule binder design than either native structures or previously designed de novo NTF2’s. Future work will focus on designing and experimentally characterizing small molecule binding proteins and enzymes using these scaffolds.

## Acknowledgement

This work was supported by a grant from the Department of Defense (DOD-0001039633, D.Z., D.B.), a grant from the National Institute on Aging (5U19AG065156, D.R.H., D.B.), a gift from the Washington Research Foundation (L.A.), a gift from Microsoft (J.D., D.B.), and, a Human Frontier Science Program Cross Disciplinary Fellowship (LT000395/2020-C, L.F.M.), an EMBO Non-Stipendiary Fellowship (ALTF 1047-2019, L.F.M.), and the Howard Hughes Medical Institute (A.C., D.B.), the Audacious Project at the Institute for Protein Design (A.K.B., A.K., L.C., D.B.), the Open Philanthropy Project Improving Protein Design Fund (B.I.M.W, H.N., D.B.), and Dr. Eric and Ms. Wendy Schmidt, and Schmidt Futures funding from Eric and Wendy Schmidt by recommendation of the Schmidt Futures program (L.C., D.B.).

This work is based upon research at the Northeastern Collaborative Access Team beamlines, which are funded by the National Institute of General Medical Sciences from the National Institutes of Health (P30 GM124165). This research used resources of the Advanced Photon Source, a U.S. Department of Energy (DOE) Office of Science User Facility operated for the DOE Office of Science by Argonne National Laboratory under Contract No. DE-AC02-06CH11357. We thank Dr. Minkyung Baek for her help on using RoseTTAFold. We thank Xinting Li for her help on verifying protein expression using electrospray ionization high-resolution mass spectroscopy. We thank Dr. Robert Ragotte for his suggestions on crystallization studies. We thank Dr. Will Sheffler for his homogeneous transform package, ‘homog’. We thank Dr. Stacey Gerben for her suggestion on plotting pocket characteristics. We thank Dr. Ivan Anishchenko for his help on sequence similarity comparison to native proteins. We thank Dr. Florian Praetorius for his editing and suggestions during manuscript preparation. We thank Microsoft and AWS for generous gifts of cloud computing credits.

This research used resources of the National Energy Research Scientific Computing Center, which is supported by the Office of Science of the U.S. Department of Energy under Contract No. DE-AC02-05CH11231.

## Online Methods

### Protein generation and sequence design pipeline

Initial models were derived through AF2 prediction of randomly generated amino acid sequences. Sequence space was traversed through mutations (1-3 mutations at a time) propagated in repeat units followed by evaluation of the predicted structure of the modified sequence. Cyclic character was evaluated as helical rise near zero, and per-unit rotation near 360/*N* degrees, where *N* is the number of repeats. For each predicted model the difference of the computed values to ideal values was calculated and then rescaled logistically to a score between 0 and 1.

Closure score is a score from 0 to 1 (0 indicates perfect closure, 1 indicates no rotation within per-unit), which is a linear combination of rescaled delta rise and rescaled delta rotation. Delta rise and delta rotation are computed by extrapolating a screw axis from smoothed relative transforms between repeat units. Relative transforms between matching repeats where derived from protein backbone positions, using the Nitrogen, **ɑ**-Carbon, and Carboxyl Carbon to define the local coordinate plane for each residue position, and then deriving the homogenous relative transform between two coordinate frames. Relative transforms were averaged by averaging the quaternions corresponding to the relative transforms, and directly computing the mean translation vector from relative transforms. This smoothed transform is then used as a proxy for a rigid body transform representing the relationship of repeat units. A helical axis is derived from this transform, as well as rise along and rotation about that axis.

The ideal rise for a cyclic repeat is 0 and an ideal rotation is 360/*N* degrees, where *N* is the number of repeats. The deltas of the rise/rotation value of the predicted structure to the ideal values were rescaled logistically to give values between 0 and 1 with midpoints at some desirable delta value.

The logistic rescaling function used was:

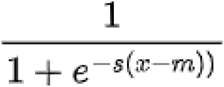

Where *s* is the slope factor, *x* is the delta value, and *m* is the logistic scale midpoint. *m* and *s* for rotation were 4 and 1.5 respectively. Those values for rise were 2 and 2, respectively. The mean between these rescaled values was used as a score for closure quality.

After generation of initial pseudocycle scaffolds, we used ProteinMPNN (Justas 2022 unpublished) to design the sequences as asymmetric monomeric proteins and used AF2 and RF to generate structural models for the new sequences.

Visual inspection of the ProteinMPNN designed pseudocycle models suggested that many designed proteins had large areas of surface apolar residues (see **supplementary Figure. S2 and S3**). To resolve this issue, we used the Rosetta sequence design suite^14^ to perform a redesign of the pseudocycles of their surface residues. For each pseudocycle model, we generated 100 sequences using ProteinMPNN (Justas 2022 unpublished) and used these newly generated sequences to generate a scaffold-specific Position Specific Score Matrix (PSSM) file. Using Rosetta sequence design (FastDesign), surface residues of the pseudocycle models with high spatial aggregation propensity^17^ (SAP) score were selected, designed with the non-hydrophobic amino acid preference provided by the PSSM file, and scored with Rosetta metrics. These sequences were then used to predict new pseudocycle structures with AF2 and RF. We chose the AF2 rank1 model (model with highest plDDT of five available AF2 models) as the final model to use. We collected AF2 metrics including plDDT and pTM along with RF metrics including plDDT, CCE, KL divergence, and RMSD of the predicted structure to the original design model (Figure. S4).

To generate the final pseudocycle list for further design applications, we examined Rosetta metrics, AF2/RF metrics, and performed some visual inspection. We removed the models which were predicted to fold into scaffolds with Ca-RMSD over 2 Å to the original model by AF2 and generated a finalized pseudocycle list which consists of 21,021 designed proteins.

### TMalign method

We used pyrosetta^18^ to calculate the average TMscore between two input proteins by averaging the TMscore obtained when each of the two pdbs was treated as the reference pdb for sequence length normalization.

### TMalign to natives

We curated a set of sequence nonredundant (using mmseq^19^), high resolution (< 1.8 Å resolution), structures from the PDB which yielded 6,111 structures. We then ran TMalign as previously described to find designs with similar native structures.

### mTMalign of 96 characterized designs against PDB

We used the mTMalign server (http://yanglab.nankai.edu.cn/mTM-align/) to compare the designs we characterized to the whole PDB, selecting the top database hit for TMalign comparison.

### Protein clustering

Structural clusters were chosen through a combination of hierarchical subdivision by DSSP character, followed by AgglomerativeCluster (from scikitlearn^20^) on an all-by-all matrix of TMalign scores. Cluster count was selected by polling of various sizes: final cluster number was chosen by minimizing the proportion of clusters with fewer than 2 members (singletons) with respect to the joint constraints of maintaining a high mean TMScore within each cluster and a low standard deviation of intra-cluster TMScores. The final clustering yielded a mean intra-group TMScore of 0.88 (not including singleton clusters) with only 9.42% singleton clusters.

### Ligand docking to pseudocycles, NTF2, and Native proteins

The pocket residues of pseudocycles were annotated using a python script which identifies the largest internal cavity bounded by the protein after converting the protein to polyalanine and then identifies all side chain residues contacting this internal cavity. The scaffold and annotation of pocket residues of NTF2s were reported previously^13^. Previously verified native small molecule-binding proteins were taken from the PDBBind database^12^. Only single-chain native small molecule-binding proteins were selected to be comparable with pseudocycles. The binding pockets were selected based on the annotation provided by PDBBind database. The native proteins were relaxed with backbone and sidechain constraints using Rosetta to remove clashes before any further computational experiments.

All 19 ligands were docked to pseudocycles, NTF2, and native proteins following the same procedure. One to eight rotamers of each ligand were extracted from PDB or Cambridge Structure Database^21^. Hydrogens were added to ligand rotamers using OpenBabel^22^ or VMD^23^ with visual inspection. The conjugation and charge were edited/added with VMD or Chimera^24^ with visual inspection. The parameter file of the ligand was generated using the python script from Rosetta application.

Rifgen/RIFdock suite^10^ was used to perform ligand docking to all the proteins. Various amino acid rotamers (referred to as RIF, or Rotamer Interaction Field) which provide hypothetical polar, aromatic, and apolar interactions to the ligand rotamer were generated for each ligand using Rifgen function with the requirements of polar interactions to all heavy atoms from the ligand rotamer. The polar interaction requirement was dropped if the generated RIF size was smaller than 1 MB, since RIFs with this degree of sparseness often do not yield meaningful docking data. With the RIF generated for each ligand, these RIFs, which encode geometry and energy information of potential interaction between amino acid rotamers and the ligand rotamers, were docked to pseudocycles, NTF2s, and native proteins at their annotated pocket residues using RIFdock function. All remaining requirements to make polar interactions to ligand heavy atoms were kept during the docking procedure. A maximum of 30 docks were allowed to be generated for each protein scaffold. Many of the protein scaffolds fail at this step to have its pocket accommodating the ligand rotamer or provide positions to hold the required interacting amino acid rotamers.

The generated docks were designed using Rosetta sequence design suite to provide score terms to identify the protein scaffolds most suitable for holding each ligand. Each generated dock was designed using a fast version of fix-backbone sequence design procedure^11^. Previous studies suggested this fast version of fix-backbone sequence design procedure generates interface metrics which are highly correlated scores generated with slow version of fix-backbone sequence design procedure, thus can be used to design high quantities of docks for selection of promising docks for binder design^11^. Interface metrics including, “contact_molecular_surface”, “ddG” were used to select the top percentile of binders designed for each ligand^11^. The top 1% of the selected docks were identified with their scaffold type origin to compare which scaffold group (Figure. 4b-c, pseudocycle, NTF2, native protein) provided the widest accommodation of different ligands.

### Expression and purification of selected proteins

Designs were reverse translated into DNA using a custom python script that attempts to maximize host-specific codon adaptation index^25^ and IDT synthesizability, which includes optimizing whole gene and local GC content as well as removing repetitive sequences, and ordered as Eblocks from IDT. Eblocks were cloned into a pET29b-derived vector with C-terminal SNAC cleavable His Tags using Golden Gate assembly and transformed into *E. coli* BL21 strain. The solubility of the proteins was first assessed using small-scale expression. 1 mL cultures were grown in a round-bottom 96 deep-well plate covered with breathable film and shaken at 200 x rpm overnight at room temperature, the cultures were harvested using centrifugation for 10 min at 4,000 x rpm and resuspended in bugbuster lysis buffer (1x bugbuster (Millipore), 25mM Tris, 100 mM NaCl, pH 8). The lysed cells were spun down and 10 uL of clear supernatant was checked for each protein on SDS-PAGE gel. Protein bands at expected molecular range were used as judgment for protein expression and solubility. Soluble designs were subsequently grown in 50 mL autoinduction media in 250 mL baffled erlenmeyer flasks for assay-scale production (6 hours at 37 ℃ followed by 24 hours at 18 ℃ shaking at 180 rpm). Cells for each design culture were harvested and resuspended in 30 mL of lysis buffer (25mM Tris 100 mM NaCl, pH 8, with protease inhibitor tablet) and sonicated to lyse (3 min sonication, 10 s pulse, 10 s pause, 60% amplitude). After centrifugation for 30 min at 14,000 x rpm, soluble fractions were bound to 1 mL Ni-NTA resin (Qiagen) in a Econo-Pac® gravity column (BIO-RAD) at 4 ℃ for 1 hour with rotation. The resin was washed with 20 CV (column volume) low salt buffer (50 mM tris, 100 mM NaCl, 50 mM Imidazole, pH 8) and with 20 CV high salt buffer (50 mM tris, 1000 mM NaCl, 50 mM Imidazole, pH 8).

For initial characterization using SEC (protocol 1) and CD, proteins were eluted with 2 CV of elution buffer (25 mM tris, 100 mM NaCl, 500 mM Imidazole, pH 8) and purified on a superdex 75 increase 10/300 GL column connected to ÄKTA protein purification systems in TBS buffer (25 mM Tris, 100 mM NaCl, pH 8).

For crystallography (protocol 2), the samples were treated identically except 4 to 8 flasks of 50 mL cultures were pooled together before sonication and His tags were cleaved on bead following the SNAC cleavage protocol^26^ before subsequent SEC purification.

### Circular dichroism characterization of selected proteins

Circular dichroism spectra were measured with a Jasco J-1500 CD spectrometer. Samples were typically around 0.25 mg/mL (range 0.1 - 0.5 mg/ml) in 25 mM phosphate buffer pH 8, and a 1-mm path length cuvette was used. The CD signal was converted to mean residue ellipticity by dividing the raw spectra by N × C × L × 10, where N is the number of residues, C is the concentration of protein, and L is the path length (0.1 cm).

### Crystallographic analysis

Crystals were produced using the sitting drop vapor diffusion method. Drops with volumes of 200 nL in ratios of 1:1, 2:1, and 1:2 protein to crystallization were set up in 96 well plates at 20°C, using the Mosquito from SPT Labtech. Drops were monitored using the JANSi UVEX imaging system.

For H10, diffraction quality crystals appeared in a mixture of 0.12 M D-glucose, 0.12 M D-mannose, 0.12 M D-galactose, 0.12 M L-fucose, 0.12 M D-xylose, 0.12 M

N-acetyl-D-glucosamine, 0.0499 M HEPES, 0.0501 M MOPS (acid), 20% v/v PEG 500 MME, and 10% w/v PEG 20,000.

For H12, diffraction quality crystals appeared in a mixture of 0.09 M sodium fluoride, 0.09 M sodium bromide, 0.09 sodium iodide, 0.0499 M HEPES, 0.0501 M MOPS (acid), 12.5% v/v MPD, 12.5% PEG 1000, and 12.5% w/v PEG 3350.

Crystals were cryoprotected prior to flash freezing in liquid nitrogen before shipping for data collection at synchrotron. Data collection was performed with synchrotron radiation at the Advanced Photon Source (APS) on beamline 24ID-C.

X-ray intensities and data reduction were evaluated and integrated using either XDS^27^ or HKL3000^28^ and merged/scaled using Pointless/Aimless in the CCP4 program suite^29^. Structure determination and refinement starting phases were obtained by molecular replacement using Phaser^30^ using the design model for the structures. Following molecular replacement, the models were improved using phenix autobuild^31^; efforts were made to reduce model bias by setting rebuild-in-place to false, and using simulated annealing. Structures were refined in Phenix^31^. Model building was performed using COOT^32^. The final model was evaluated using MolProbity^33^. Data collection and refinement statistics are recorded in supplementary Table S3. Data deposition, atomic coordinates, and structure factors reported in this paper will be deposited in the PDB (PDB ID XXXX, YYYY).

## Supporting Information

**Figure S1.**
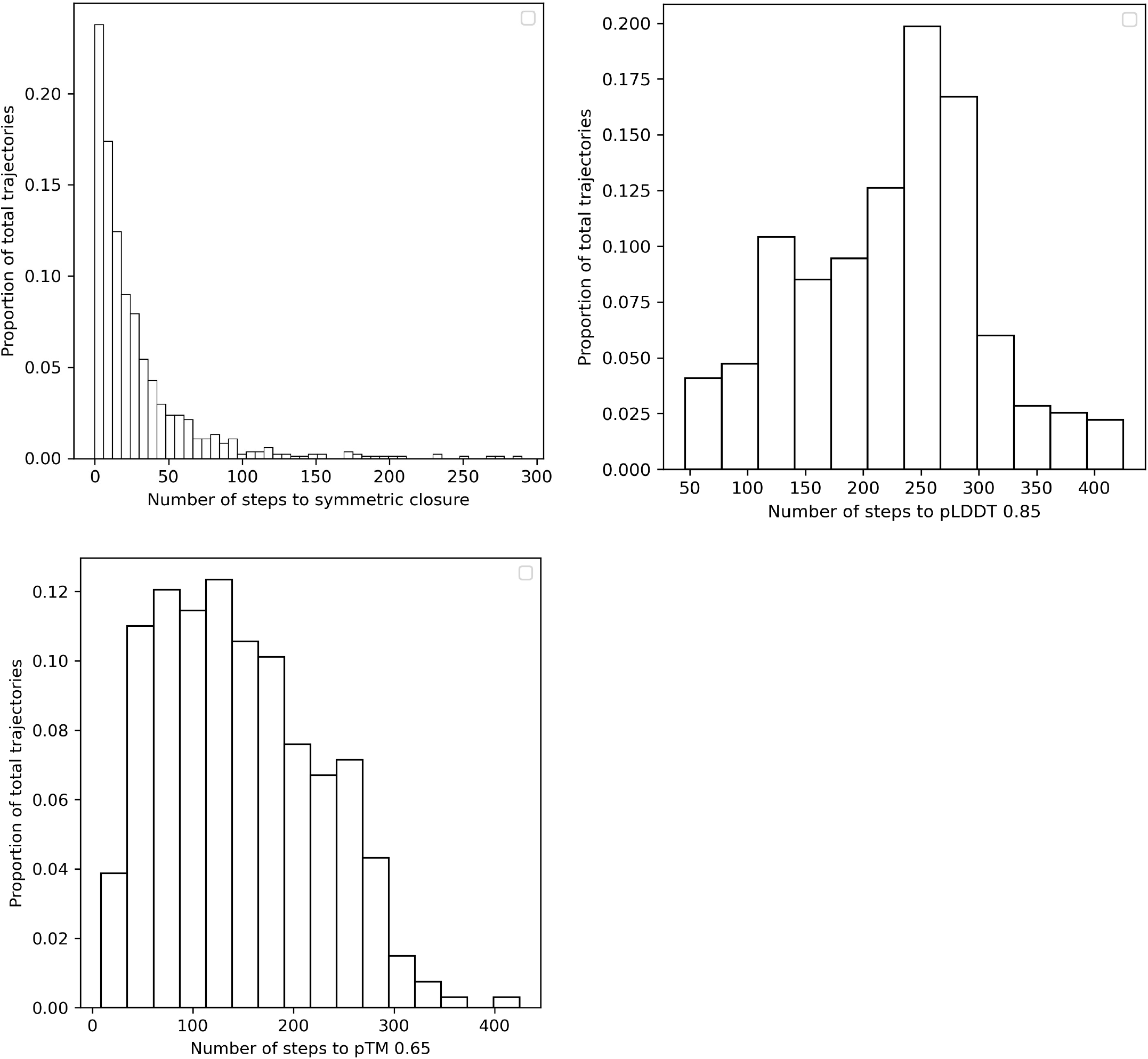
The histograms of MCMC steps to convergence. The number of steps to closure for a representative sample of trajectories is shown here. Symmetric closure is defined as a “closure score” of 0.1 or less. A clear trend is that AF2 readily predicts closed, cyclic structures from random repetitive sequences, but with very low confidence and quality until a rather large number of mutations.

**Figure S2.**
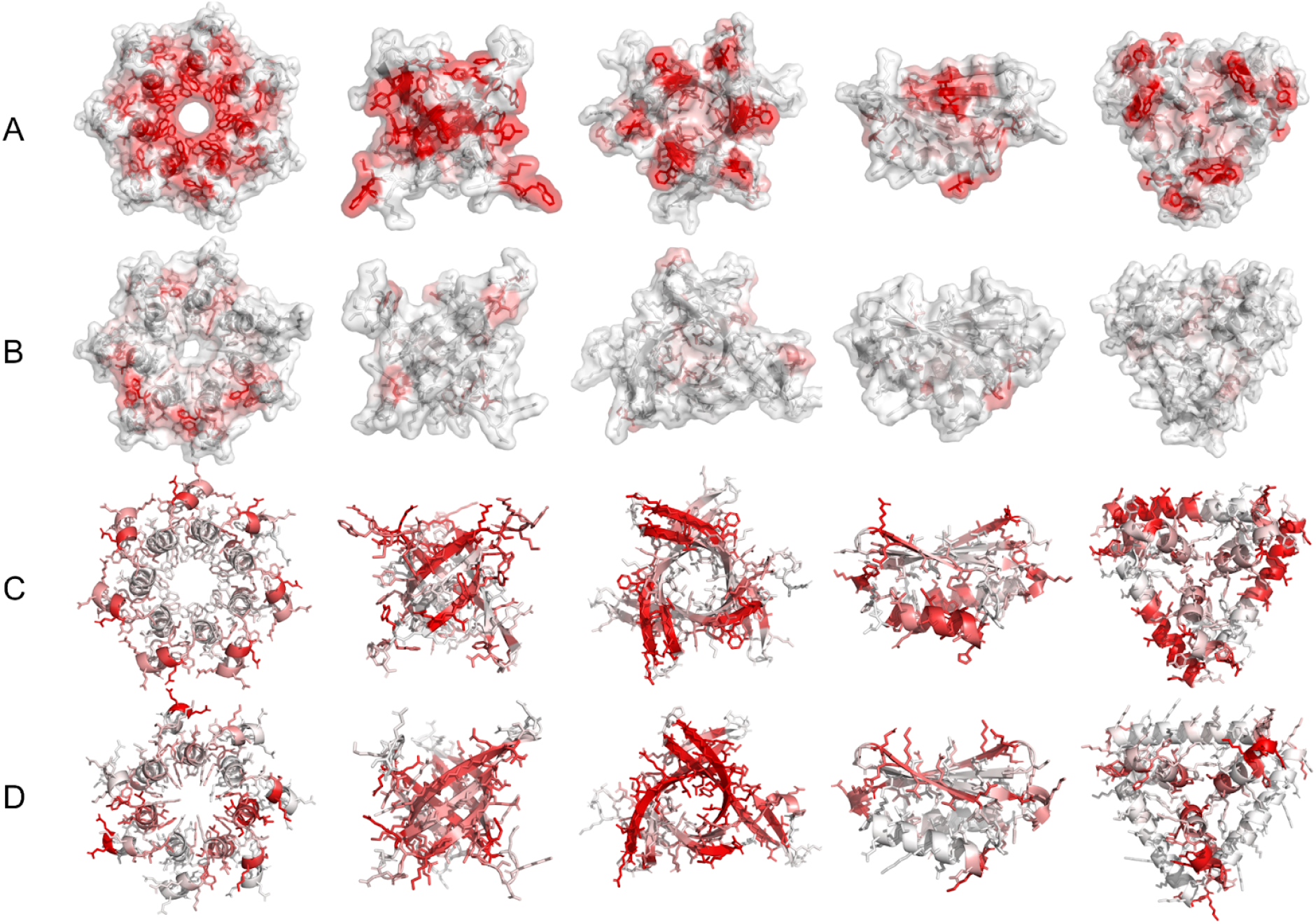
Cartoons of exemplary designed pseudocycles from cyclic repeat protein hallucination showing per residue SAP and psipred scores before and after ProteinMPNN+Rosetta redesign. a) 5 diverse representative proteins following the hallucination procedure colored by SAP score. Color scales from white (no aggregation propensity) to red (high aggregate propensity). b) The same 5 proteins after ProteinMPNN redesign and Rosetta surface optimization colored by SAP score. c) The same 5 hallucinated proteins colored by agreement of single sequence psipred prediction with the intended secondary structure. Color scales from white (perfect agreement) to red (no agreement). d) The redesigned proteins colored by agreement of single sequence psipred prediction with the intended secondary structure.

**Figure S3.**
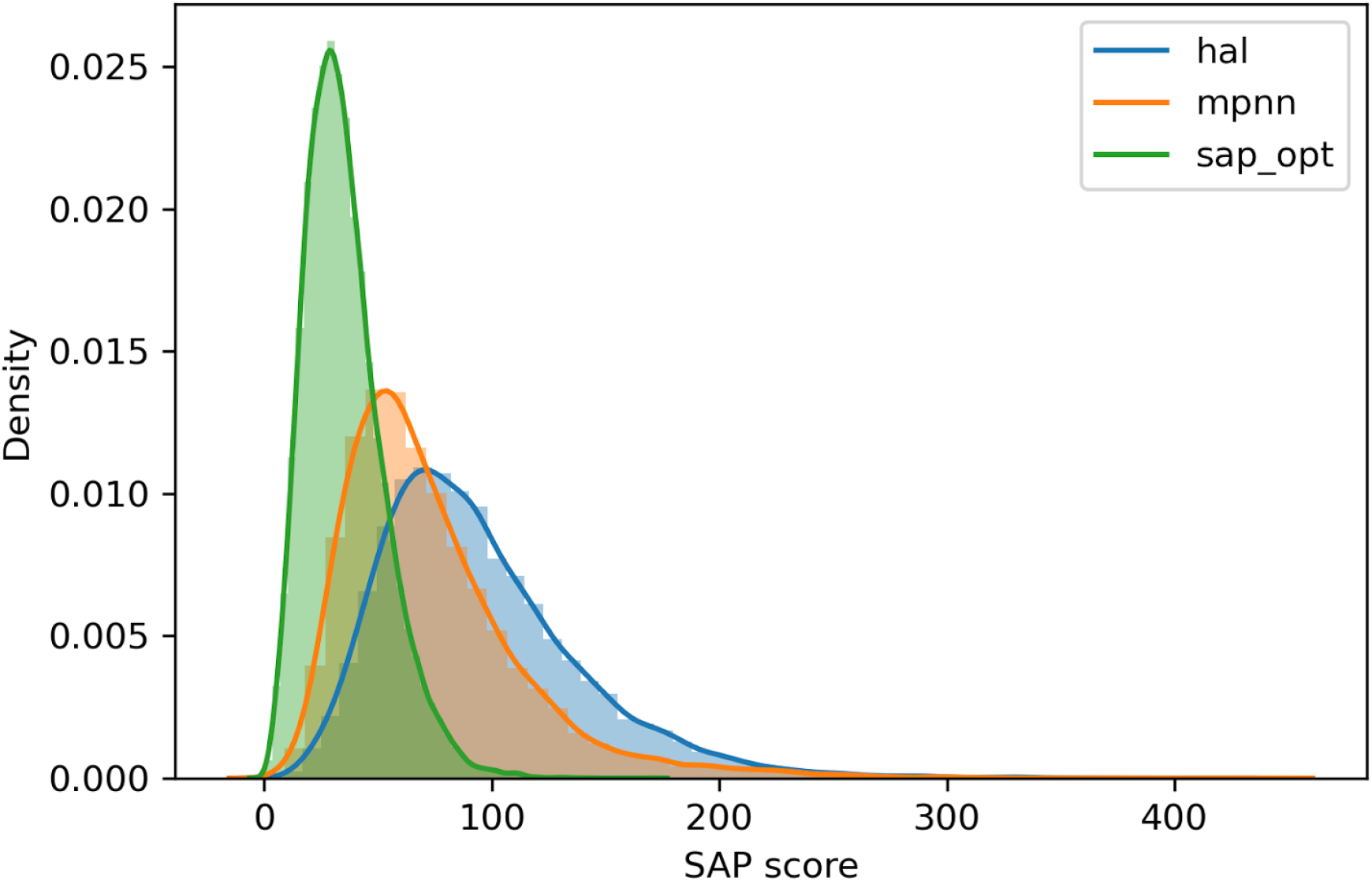
SAP score improvement during design. Histogram of SAP score for original hallucinations (hal), after ProteinMPNN (mpnn) redesign, and after Rosetta surface optimization (sap_opt).

**Figure S4.**
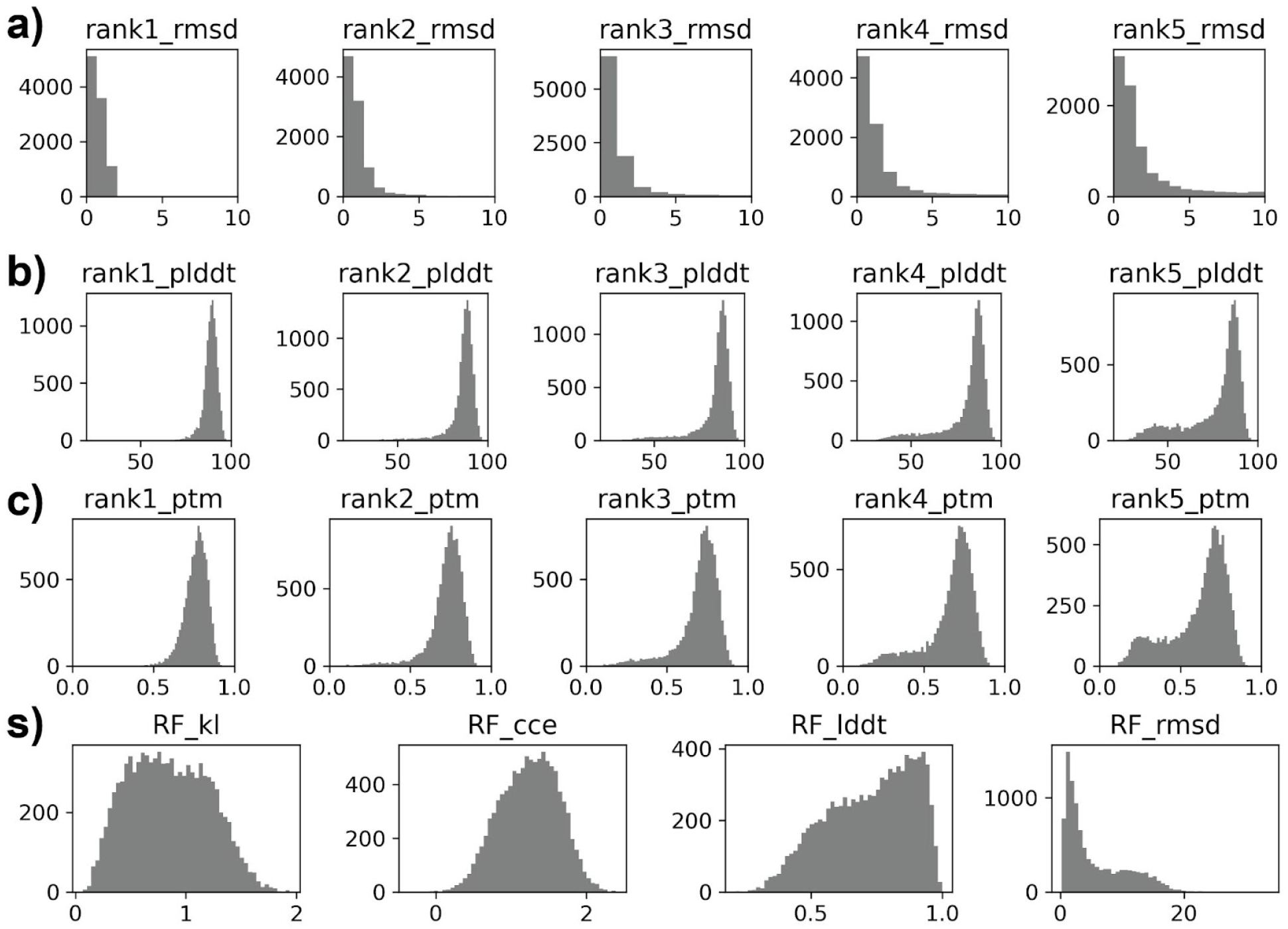
AF2 and RF metric histograms for 9838 pseudocycle cluster representatives. a). AF2 Ca-RMSD to design models for 5 AF2 models by AF2 rank. b). AF2 plDDT for predictions. c). AF2 ptm for predictions. d). RosettaFold (RF) Ca-RMSD to design model, RF lddt, RF KL divergence, and RF CCE for predictions.

**Figure S5.**
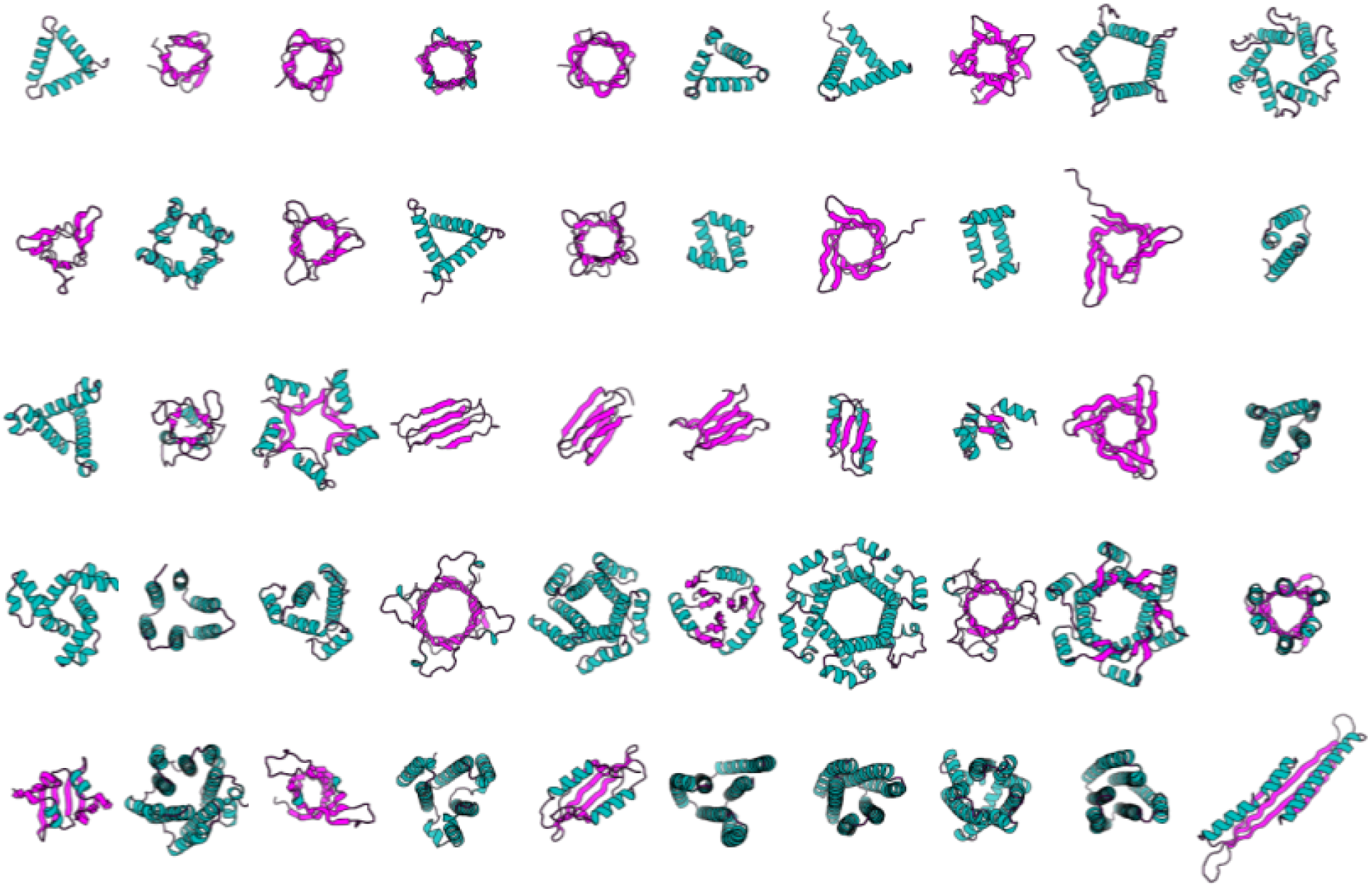
50 Diverse pseudocycle structures.

**Figure S6.**
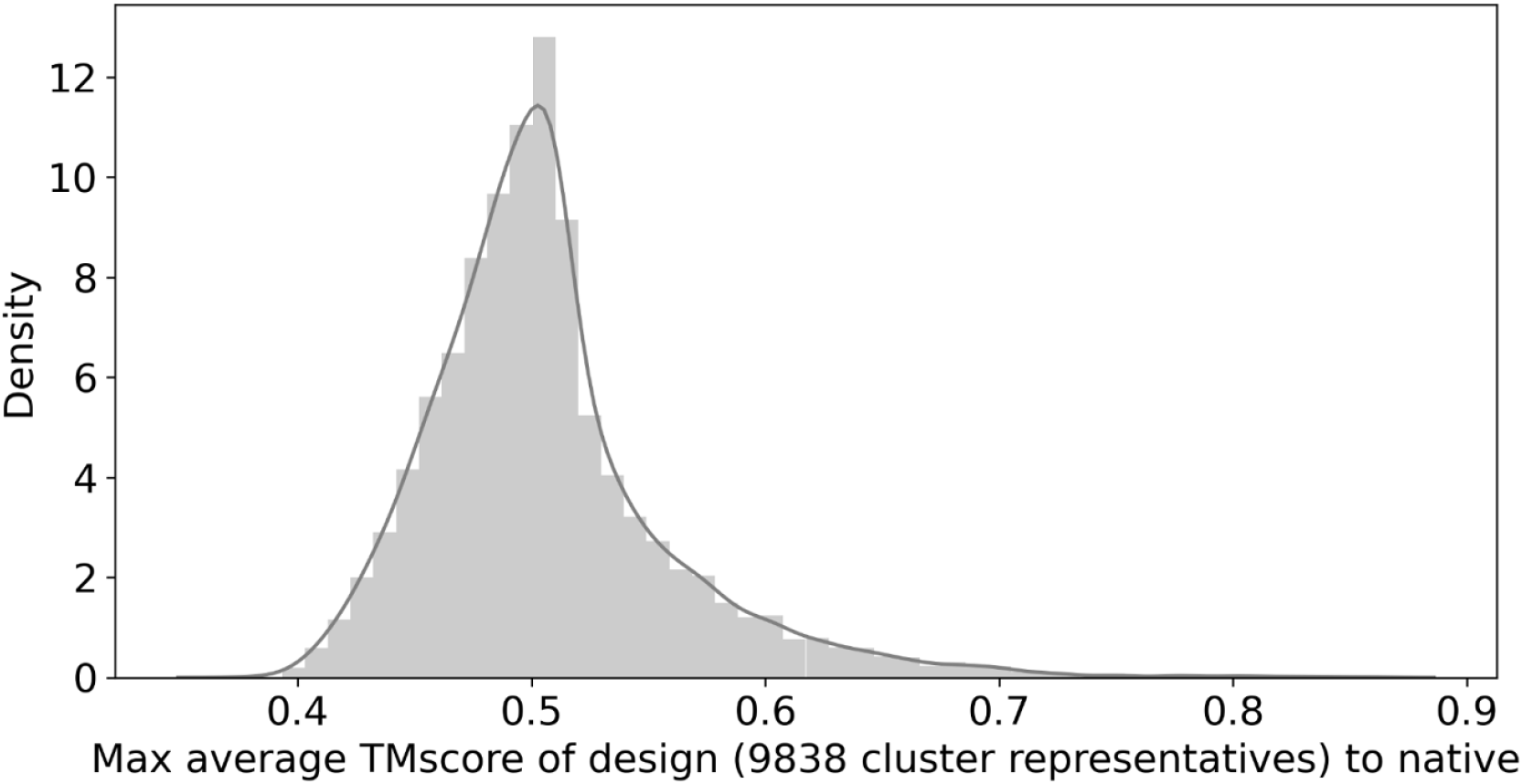
TMscore of designs to natives. For each of our 9838 design cluster representatives, we computed the max average TMscore to native structures and plotted as a histogram.

**Figure S7.**
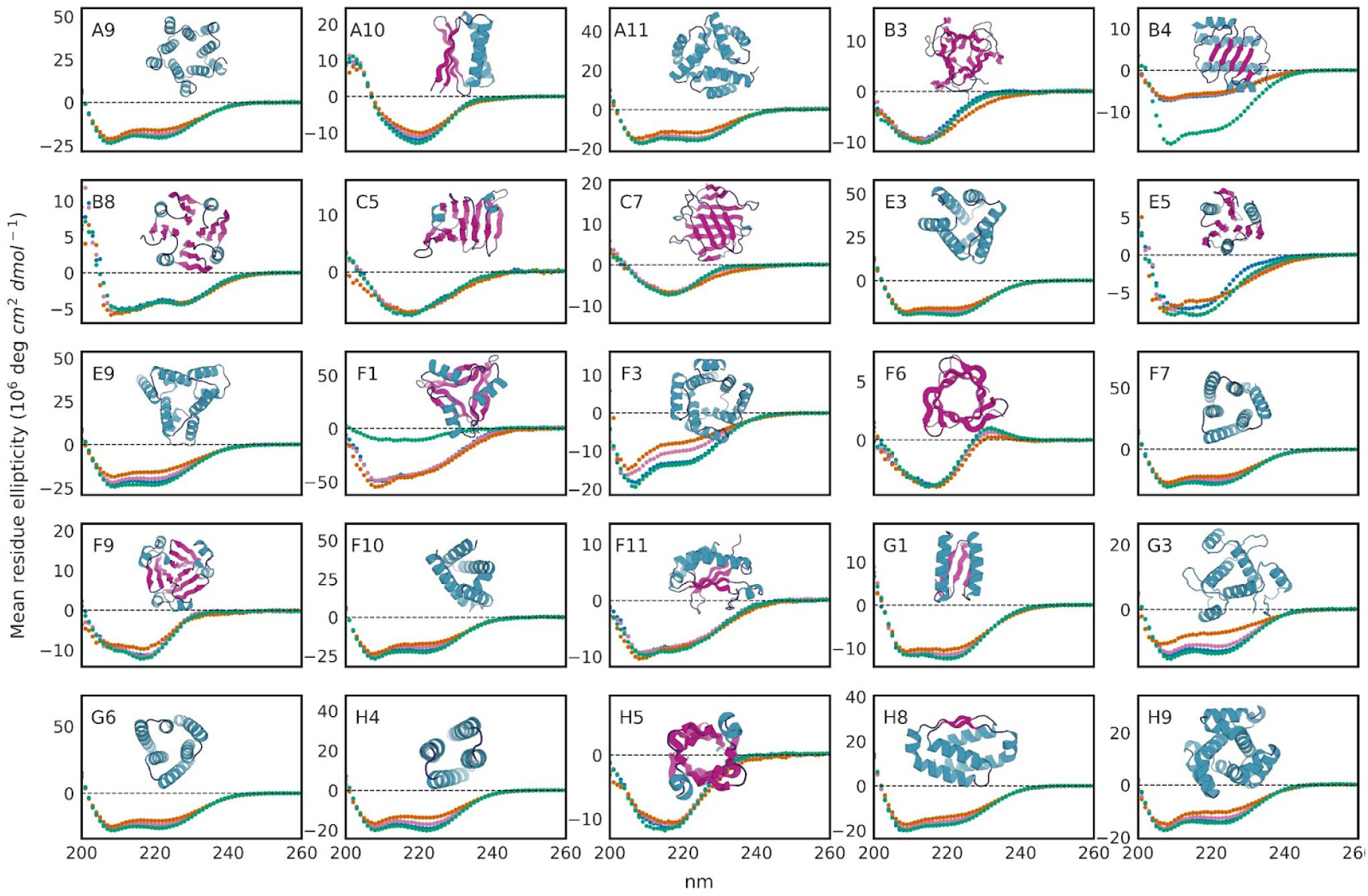
CD data for 25 designs not shown in figure 2. Different temperatures of the CD scan spectra are plotted as follows: 25 °C in blue, 55 °C in orange, 95 °C in pink, refolding at 25 °C in green. The cartoon of the corresponding designed pseudocycle is shown with each CD spectra. The sheet, helix, loop substructures are colored in magenta, teal, and dark blue, respectively.

**Figure S8.**
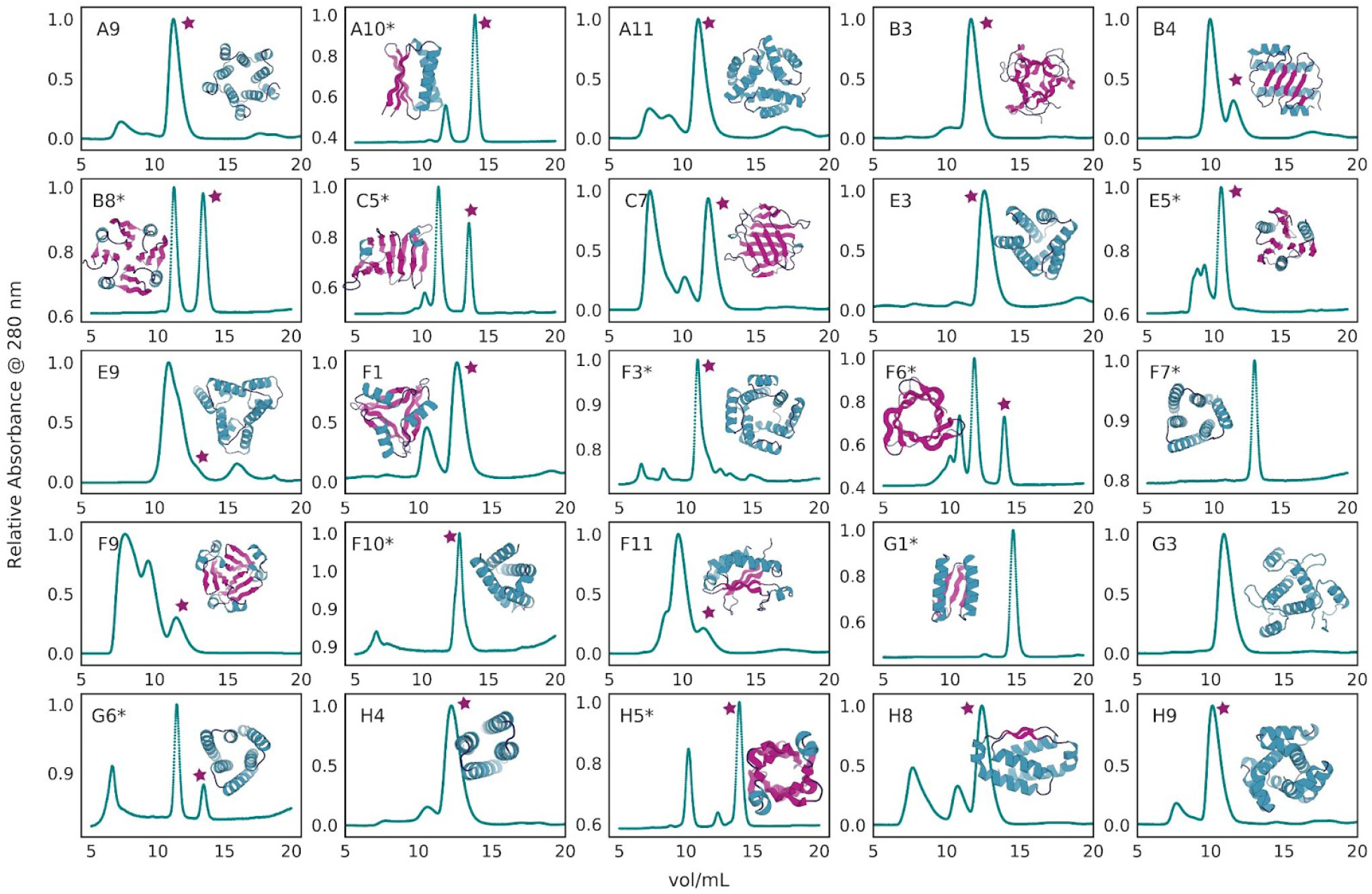
SEC data for 25 designs not shown in figure 2. The SEC analysis performed using protocol2 were marked with a star (*) at its label. Monomeric fraction was marked out using a magenta star. The cartoon of the corresponding designed pseudocycle is shown with each subplot. The sheet, helix, loop substructures are colored in magenta, teal, and dark blue, respectively.

**Figure S9.**
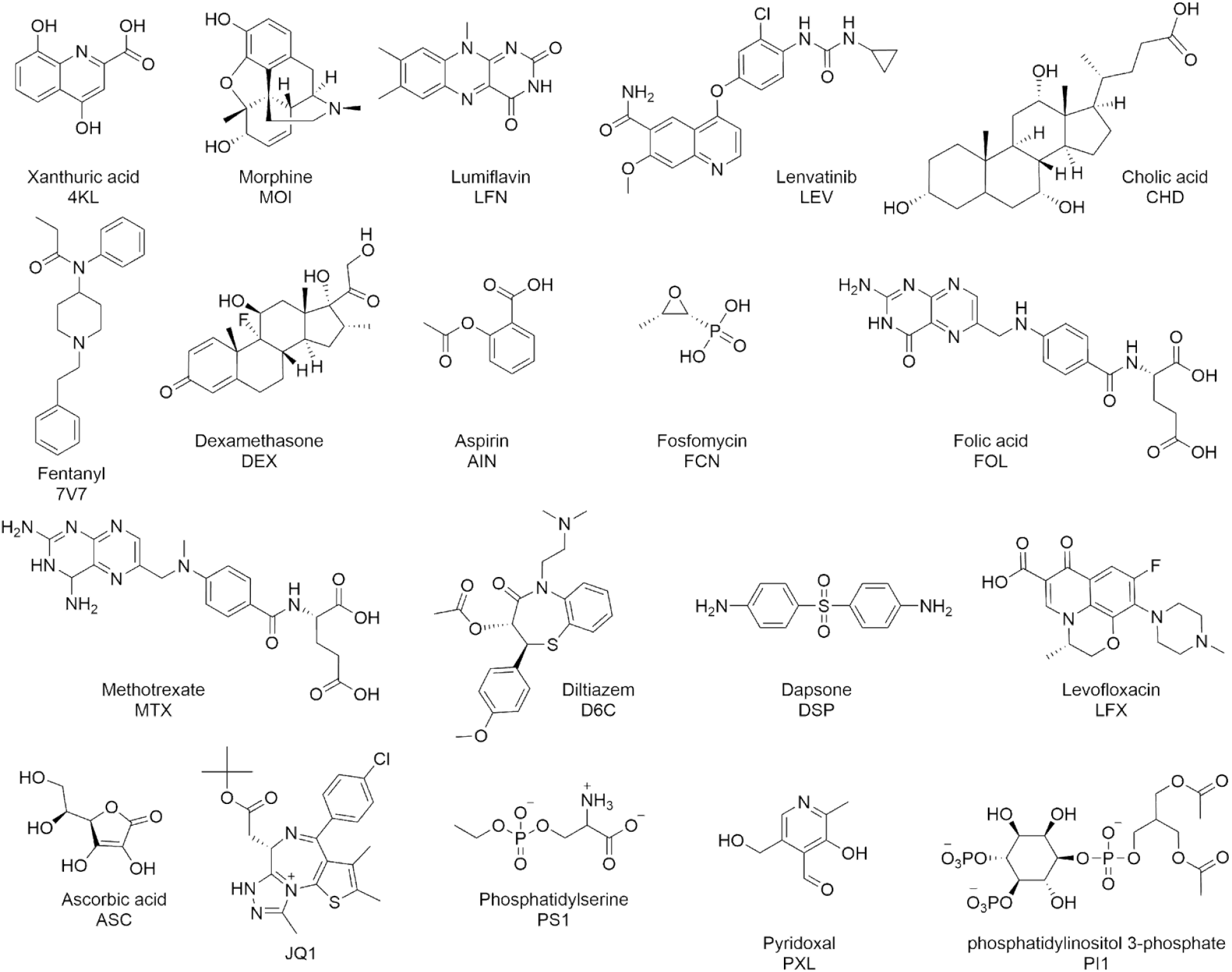
Ligand structures used for docking. The structure of ligands used for docking and design in this study. The three-letter names are marked out with the chemical name and structures.

**Figure S10.**
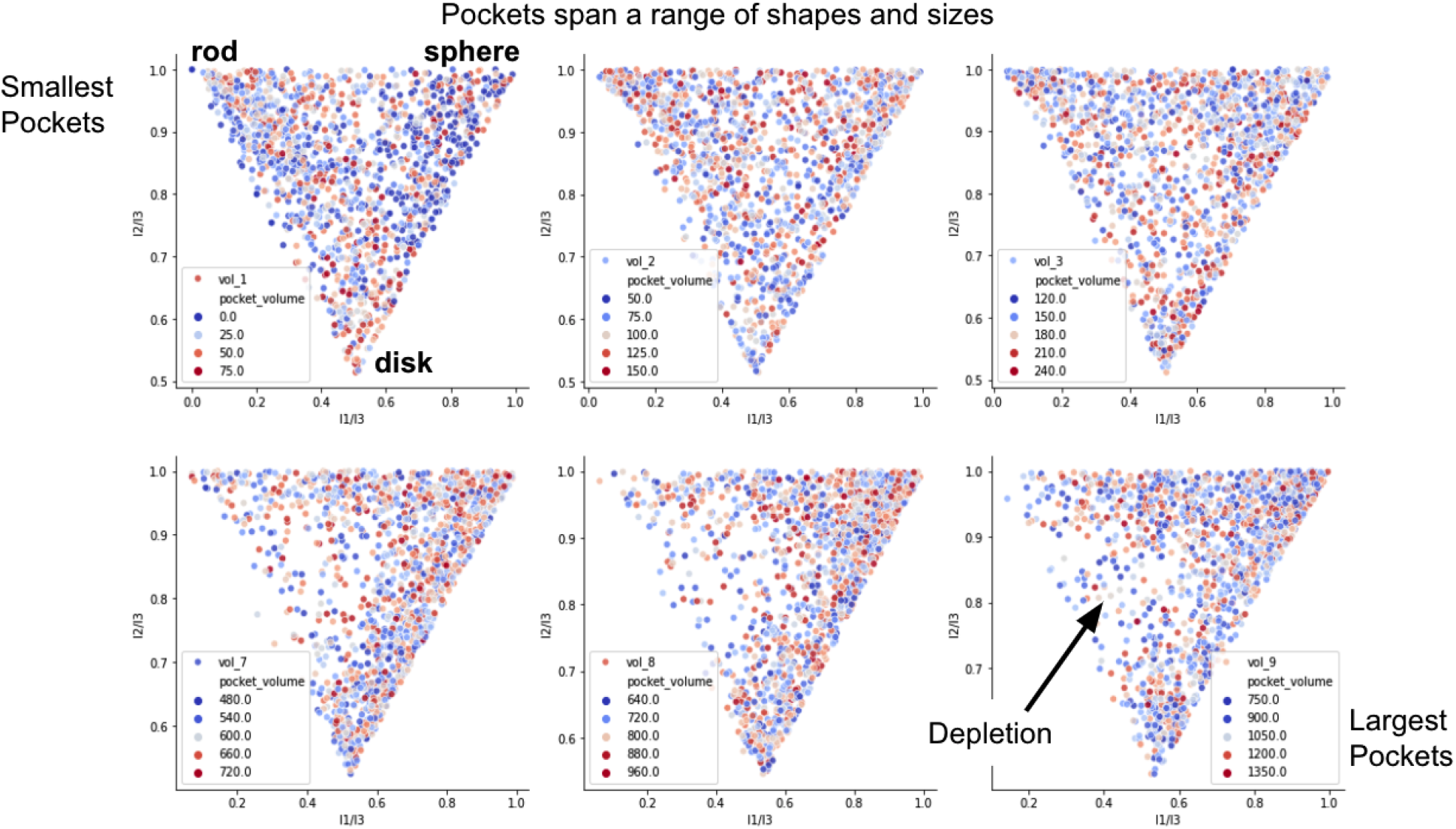
pocket shapes. Pockets were detected for 21,021 designs with sequences converted to poly-alanine to show max possible pocket size. Plots show the I2/I3 vs I1/I3 ratio which dictate the pocket shape. We show plots for pockets binned from small (75 cubic Å) to large (1400 cubic Å).

**Table S1.**
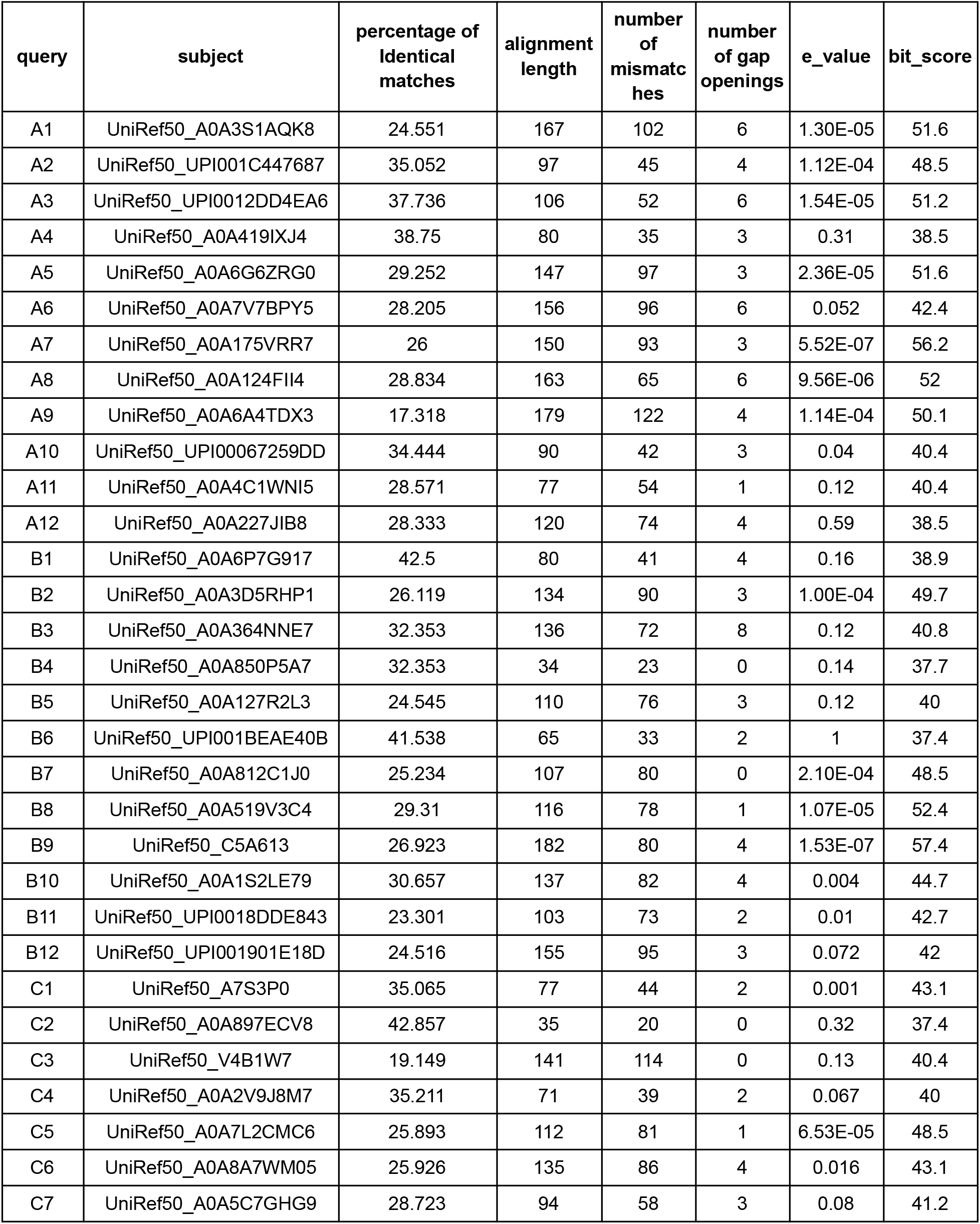
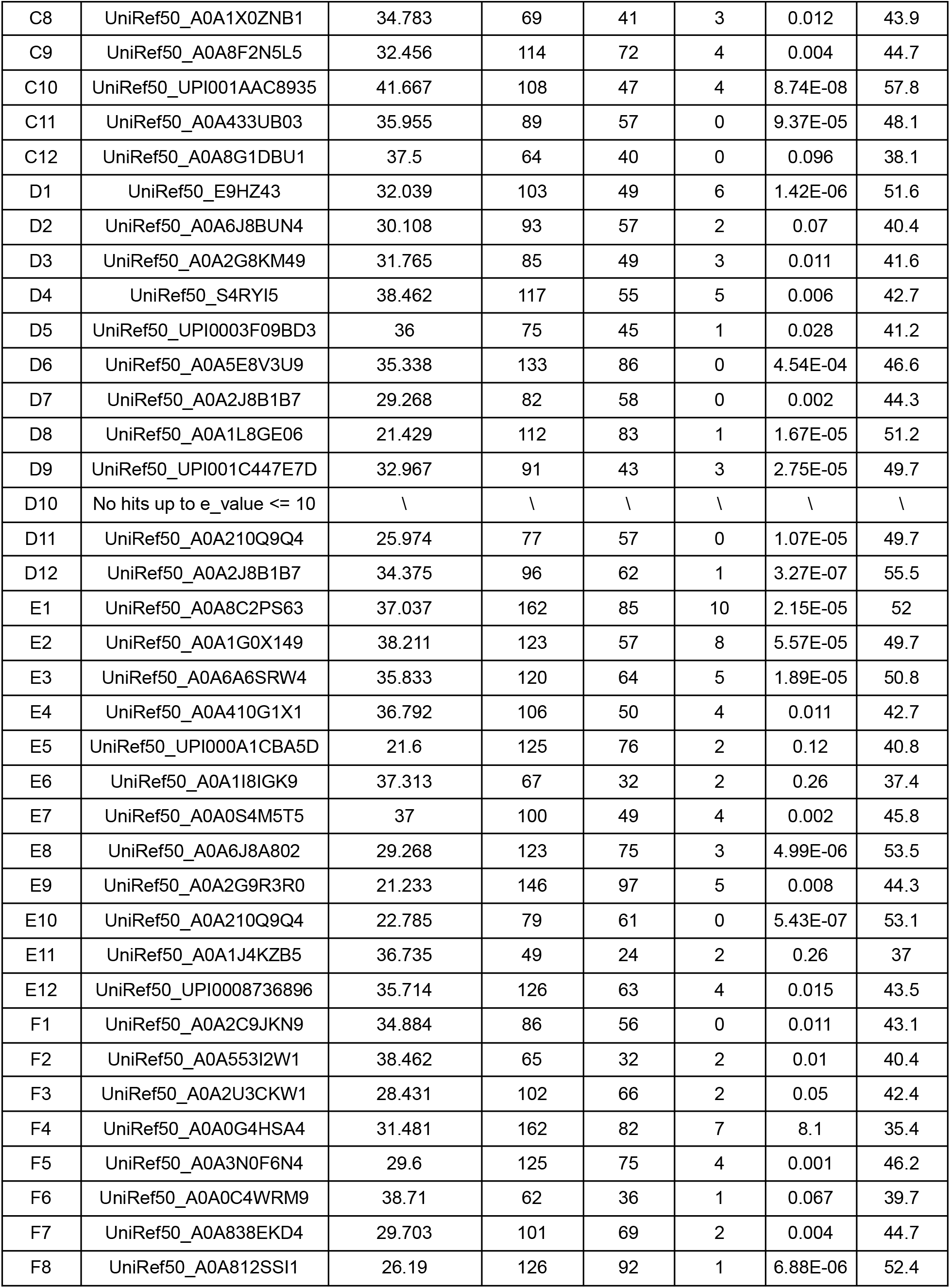
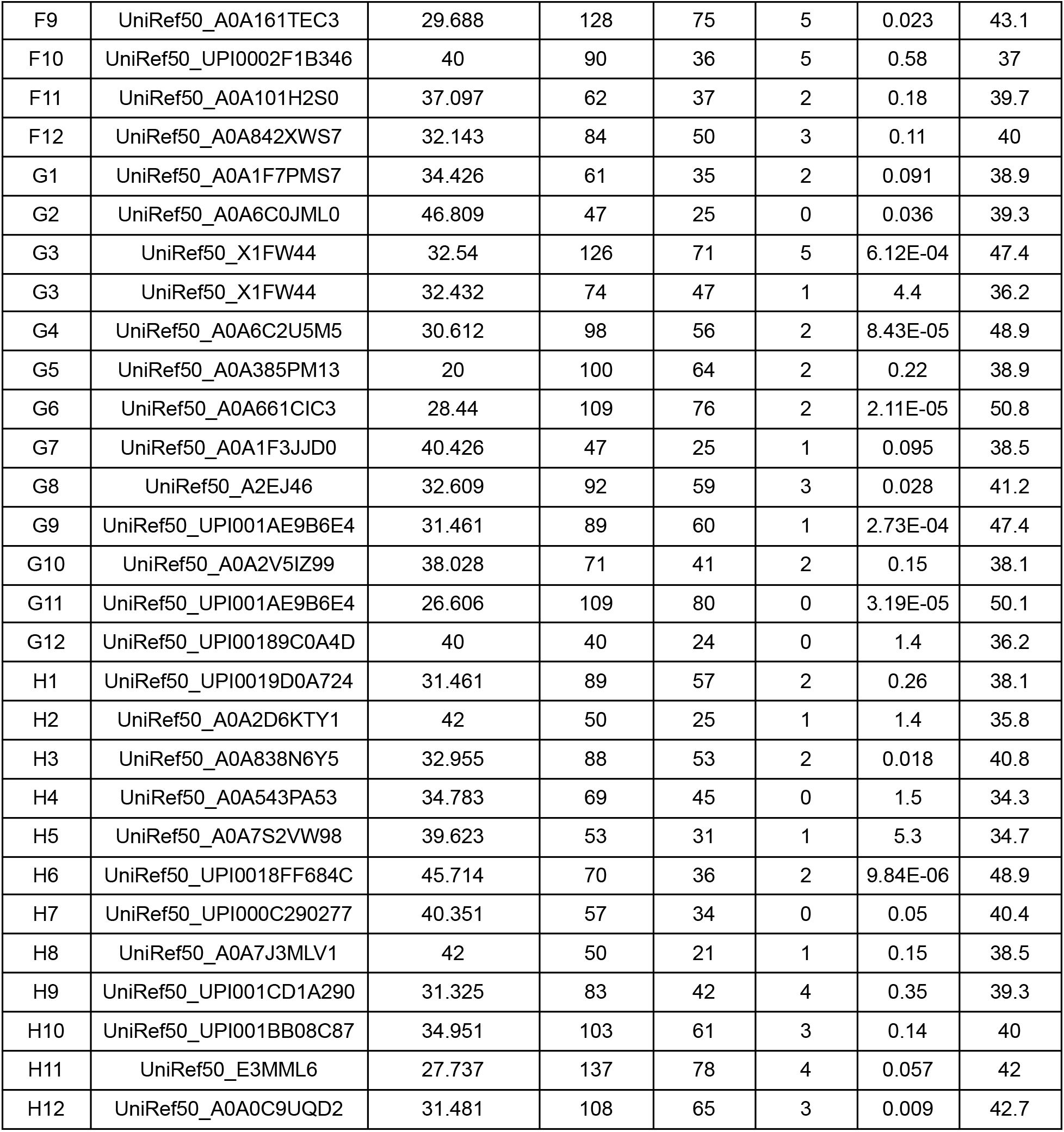
The sequence similarity of 96 experimentally-characterized pseudocycles to native proteins (UniRef50).

| query | subject | percentage of identical matches | alignment length | number of mismatches | number of gap openings | e_value | bit_score |
| --- | --- | --- | --- | --- | --- | --- | --- |
| A1 | UniRef50_A0A3S1AQK8 | 24.551 | 167 | 102 | 6 | 1.30E-05 | 51.6 |
| A2 | UniRef50_UPI001C447687 | 35.052 | 97 | 45 | 4 | 1.12E-04 | 48.5 |
| A3 | UniRef50_UPI0012DD4EA6 | 37.736 | 106 | 52 | 6 | 1.54E-05 | 51.2 |
| A4 | UniRef50_A0A419IXJ4 | 38.75 | 80 | 35 | 3 | 0.31 | 38.5 |
| A5 | UniRef50_A0A6G6ZRG0 | 29.252 | 147 | 97 | 3 | 2.36E-05 | 51.6 |
| A6 | UniRef50_A0A7V7BPY5 | 28.205 | 156 | 96 | 6 | 0.052 | 42.4 |
| A7 | UniRef50_A0A175VRR7 | 26 | 150 | 93 | 3 | 5.52E-07 | 56.2 |
| A8 | UniRef50_A0A124FII4 | 28.834 | 163 | 65 | 6 | 9.56E-06 | 52 |
| A9 | UniRef50_A0A6A4TDX3 | 17.318 | 179 | 122 | 4 | 1.14E-04 | 50.1 |
| A10 | UniRef50_UPI00067259DD | 34.444 | 90 | 42 | 3 | 0.04 | 40.4 |
| A11 | UniRef50_A0A4C1WNI5 | 28.571 | 77 | 54 | 1 | 0.12 | 40.4 |
| A12 | UniRef50_A0A227JIB8 | 28.333 | 120 | 74 | 4 | 0.59 | 38.5 |
| B1 | UniRef50_A0A6P7G917 | 42.5 | 80 | 41 | 4 | 0.16 | 38.9 |
| B2 | UniRef50_A0A3D5RHP1 | 26.119 | 134 | 90 | 3 | 1.00E-04 | 49.7 |
| B3 | UniRef50_A0A364NNE7 | 32.353 | 136 | 72 | 8 | 0.12 | 40.8 |
| B4 | UniRef50_A0A850P5A7 | 32.353 | 34 | 23 | 0 | 0.14 | 37.7 |
| B5 | UniRef50_A0A127R2L3 | 24.545 | 110 | 76 | 3 | 0.12 | 40 |
| B6 | UniRef50_UPI001BEAE40B | 41.538 | 65 | 33 | 2 | 1 | 37.4 |
| B7 | UniRef50_A0A812C1J0 | 25.234 | 107 | 80 | 0 | 2.10E-04 | 48.5 |
| B8 | UniRef50_A0A519V3C4 | 29.31 | 116 | 78 | 1 | 1.07E-05 | 52.4 |
| B9 | UniRef50_C5A613 | 26.923 | 182 | 80 | 4 | 1.53E-07 | 57.4 |
| B10 | UniRef50_A0A1S2LE79 | 30.657 | 137 | 82 | 4 | 0.004 | 44.7 |
| B11 | UniRef50_UPI0018DDE843 | 23.301 | 103 | 73 | 2 | 0.01 | 42.7 |
| B12 | UniRef50_UPI001901E18D | 24.516 | 155 | 95 | 3 | 0.072 | 42 |
| C1 | UniRef50_A7S3P0 | 35.065 | 77 | 44 | 2 | 0.001 | 43.1 |
| C2 | UniRef50_A0A897ECV8 | 42.857 | 35 | 20 | 0 | 0.32 | 37.4 |
| C3 | UniRef50_V4B1W7 | 19.149 | 141 | 114 | 0 | 0.13 | 40.4 |
| C4 | UniRef50_A0A2V9J8M7 | 35.211 | 71 | 39 | 2 | 0.067 | 40 |
| C5 | UniRef50_A0A7L2CMC6 | 25.893 | 112 | 81 | 1 | 6.53E-05 | 48.5 |
| C6 | UniRef50_A0A8A7WM05 | 25.926 | 135 | 86 | 4 | 0.016 | 43.1 |
| C7 | UniRef50_A0A5C7GHG9 | 28.723 | 94 | 58 | 3 | 0.08 | 41.2 |
| C8 | UniRef50_A0A1X0ZNB1 | 34.783 | 69 | 41 | 3 | 0.012 | 43.9 |
| C9 | UniRef50_A0A8F2N5L5 | 32.456 | 114 | 72 | 4 | 0.004 | 44.7 |
| C10 | UniRef50_UPI001AAC8935 | 41.667 | 108 | 47 | 4 | 8.74E-08 | 57.8 |
| C11 | UniRef50_A0A433UB03 | 35.955 | 89 | 57 | 0 | 9.37E-05 | 48.1 |
| C12 | UniRef50_A0A8G1DBU1 | 37.5 | 64 | 40 | 0 | 0.096 | 38.1 |
| D1 | UniRef50_E9HZ43 | 32.039 | 103 | 49 | 6 | 1.42E-06 | 51.6 |
| D2 | UniRef50_A0A6J8BUN4 | 30.108 | 93 | 57 | 2 | 0.07 | 40.4 |
| D3 | UniRef50_A0A2G8KM49 | 31.765 | 85 | 49 | 3 | 0.011 | 41.6 |
| D4 | UniRef50_S4RYI5 | 38.462 | 117 | 55 | 5 | 0.006 | 42.7 |
| D5 | UniRef50_UPI0003F09BD3 | 36 | 75 | 45 | 1 | 0.028 | 41.2 |
| D6 | UniRef50_A0A5E8V3U9 | 35.338 | 133 | 86 | 0 | 4.54E-04 | 46.6 |
| D7 | UniRef50_A0A2J8B1B7 | 29.268 | 82 | 58 | 0 | 0.002 | 44.3 |
| D8 | UniRef50_A0A1L8GE06 | 21.429 | 112 | 83 | 1 | 1.67E-05 | 51.2 |
| D9 | UniRef50_UPI001C447E7D | 32.967 | 91 | 43 | 3 | 2.75E-05 | 49.7 |
| D10 | No hits up to e_value <= 10 | \ | \ | \ | \ | \ | \ |
| D11 | UniRef50_A0A210Q9Q4 | 25.974 | 77 | 57 | 0 | 1.07E-05 | 49.7 |
| D12 | UniRef50_A0A2J8B1B7 | 34.375 | 96 | 62 | 1 | 3.27E-07 | 55.5 |
| E1 | UniRef50_A0A8C2PS63 | 37.037 | 162 | 85 | 10 | 2.15E-05 | 52 |
| E2 | UniRef50_A0A1G0X149 | 38.211 | 123 | 57 | 8 | 5.57E-05 | 49.7 |
| E3 | UniRef50_A0A6A6SRW4 | 35.833 | 120 | 64 | 5 | 1.89E-05 | 50.8 |
| E4 | UniRef50_A0A410G1X1 | 36.792 | 106 | 50 | 4 | 0.011 | 42.7 |
| E5 | UniRef50_UPI000A1CBA5D | 21.6 | 125 | 76 | 2 | 0.12 | 40.8 |
| E6 | UniRef50_A0A1I8IGK9 | 37.313 | 67 | 32 | 2 | 0.26 | 37.4 |
| E7 | UniRef50_A0A0S4M5T5 | 37 | 100 | 49 | 4 | 0.002 | 45.8 |
| E8 | UniRef50_A0A6J8A802 | 29.268 | 123 | 75 | 3 | 4.99E-06 | 53.5 |
| E9 | UniRef50_A0A2G9R3R0 | 21.233 | 146 | 97 | 5 | 0.008 | 44.3 |
| E10 | UniRef50_A0A210Q9Q4 | 22.785 | 79 | 61 | 0 | 5.43E-07 | 53.1 |
| E11 | UniRef50_A0A1J4KZB5 | 36.735 | 49 | 24 | 2 | 0.26 | 37 |
| E12 | UniRef50_UPI0008736896 | 35.714 | 126 | 63 | 4 | 0.015 | 43.5 |
| F1 | UniRef50_A0A2C9JKN9 | 34.884 | 86 | 56 | 0 | 0.011 | 43.1 |
| F2 | UniRef50_A0A553I2W1 | 38.462 | 65 | 32 | 2 | 0.01 | 40.4 |
| F3 | UniRef50_A0A2U3CKW1 | 28.431 | 102 | 66 | 2 | 0.05 | 42.4 |
| F4 | UniRef50_A0A0G4HSA4 | 31.481 | 162 | 82 | 7 | 8.1 | 35.4 |
| F5 | UniRef50_A0A3N0F6N4 | 29.6 | 125 | 75 | 4 | 0.001 | 46.2 |
| F6 | UniRef50_A0A0C4WRM9 | 38.71 | 62 | 36 | 1 | 0.067 | 39.7 |
| F7 | UniRef50_A0A838EKD4 | 29.703 | 101 | 69 | 2 | 0.004 | 44.7 |
| F8 | UniRef50_A0A812SSI1 | 26.19 | 126 | 92 | 1 | 6.88E-06 | 52.4 |
| F9 | UniRef50_A0A161TEC3 | 29.688 | 128 | 75 | 5 | 0.023 | 43.1 |
| F10 | UniRef50_UPI0002F1B346 | 40 | 90 | 36 | 5 | 0.58 | 37 |
| F11 | UniRef50_A0A101H2S0 | 37.097 | 62 | 37 | 2 | 0.18 | 39.7 |
| F12 | UniRef50_A0A842XWS7 | 32.143 | 84 | 50 | 3 | 0.11 | 40 |
| G1 | UniRef50_A0A1F7PMS7 | 34.426 | 61 | 35 | 2 | 0.091 | 38.9 |
| G2 | UniRef50_A0A6C0JML0 | 46.809 | 47 | 25 | 0 | 0.036 | 39.3 |
| G3 | UniRef50_X1FW44 | 32.54 | 126 | 71 | 5 | 6.12E-04 | 47.4 |
| G3 | UniRef50_X1FW44 | 32.432 | 74 | 47 | 1 | 4.4 | 36.2 |
| G4 | UniRef50_A0A6C2U5M5 | 30.612 | 98 | 56 | 2 | 8.43E-05 | 48.9 |
| G5 | UniRef50_A0A385PM13 | 20 | 100 | 64 | 2 | 0.22 | 38.9 |
| G6 | UniRef50_A0A661CIC3 | 28.44 | 109 | 76 | 2 | 2.11E-05 | 50.8 |
| G7 | UniRef50_A0A1F3JJD0 | 40.426 | 47 | 25 | 1 | 0.095 | 38.5 |
| G8 | UniRef50_A2EJ46 | 32.609 | 92 | 59 | 3 | 0.028 | 41.2 |
| G9 | UniRef50_UPI001AE9B6E4 | 31.461 | 89 | 60 | 1 | 2.73E-04 | 47.4 |
| G10 | UniRef50_A0A2V5IZ99 | 38.028 | 71 | 41 | 2 | 0.15 | 38.1 |
| G11 | UniRef50_UPI001AE9B6E4 | 26.606 | 109 | 80 | 0 | 3.19E-05 | 50.1 |
| G12 | UniRef50_UPI00189C0A4D | 40 | 40 | 24 | 0 | 1.4 | 36.2 |
| H1 | UniRef50_UPI0019D0A724 | 31.461 | 89 | 57 | 2 | 0.26 | 38.1 |
| H2 | UniRef50_A0A2D6KTY1 | 42 | 50 | 25 | 1 | 1.4 | 35.8 |
| H3 | UniRef50_A0A838N6Y5 | 32.955 | 88 | 53 | 2 | 0.018 | 40.8 |
| H4 | UniRef50_A0A543PA53 | 34.783 | 69 | 45 | 0 | 1.5 | 34.3 |
| H5 | UniRef50_A0A7S2VW98 | 39.623 | 53 | 31 | 1 | 5.3 | 34.7 |
| H6 | UniRef50_UPI0018FF684C | 45.714 | 70 | 36 | 2 | 9.84E-06 | 48.9 |
| H7 | UniRef50_UPI000C290277 | 40.351 | 57 | 34 | 0 | 0.05 | 40.4 |
| H8 | UniRef50_A0A7J3MLV1 | 42 | 50 | 21 | 1 | 0.15 | 38.5 |
| H9 | UniRef50_UPI001CD1A290 | 31.325 | 83 | 42 | 4 | 0.35 | 39.3 |
| H10 | UniRef50_UPI001BB08C87 | 34.951 | 103 | 61 | 3 | 0.14 | 40 |
| H11 | UniRef50_E3MML6 | 27.737 | 137 | 78 | 4 | 0.057 | 42 |
| H12 | UniRef50_A0A0C9UQD2 | 31.481 | 108 | 65 | 3 | 0.009 | 42.7 |

**Table S2.**
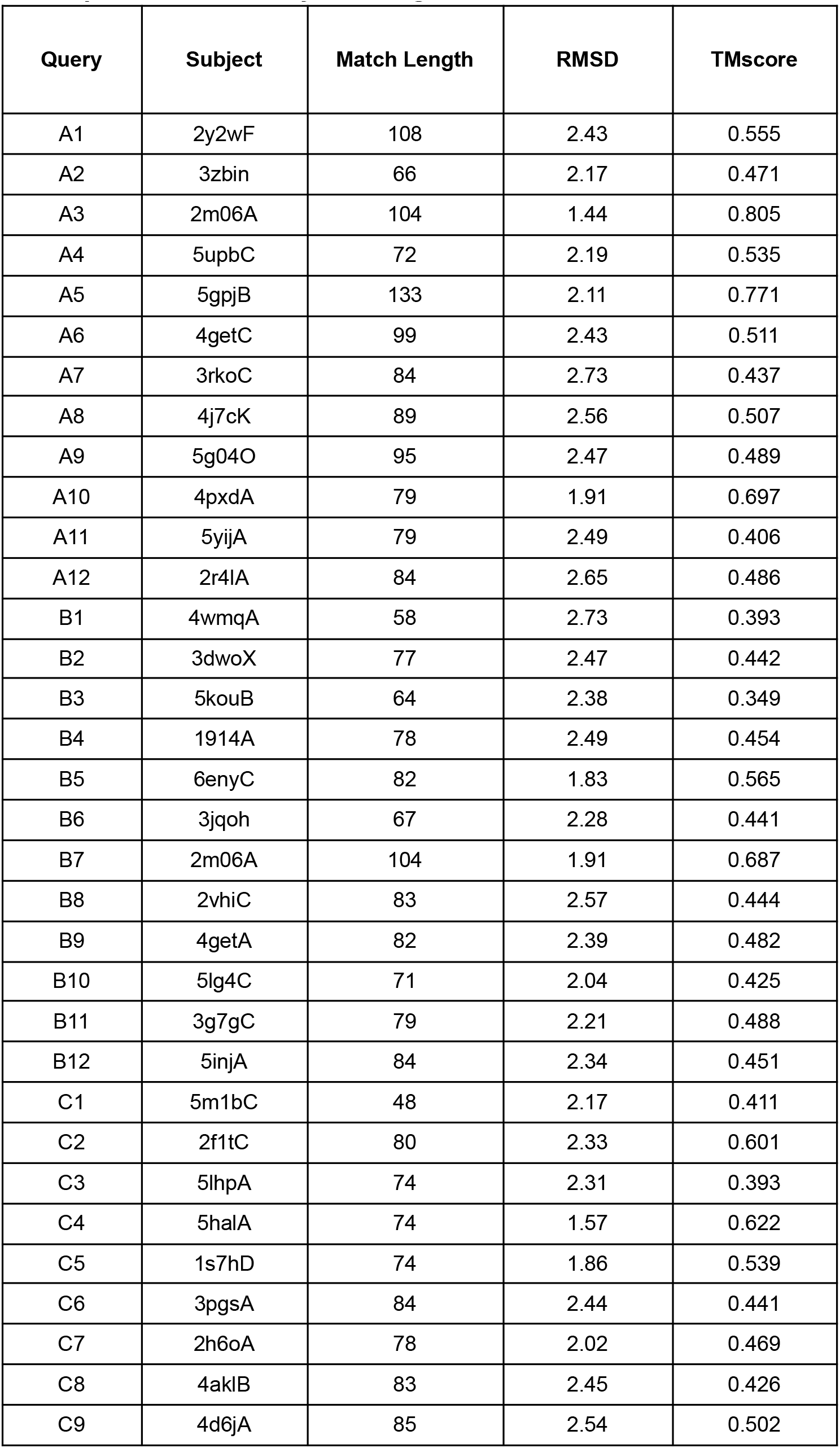
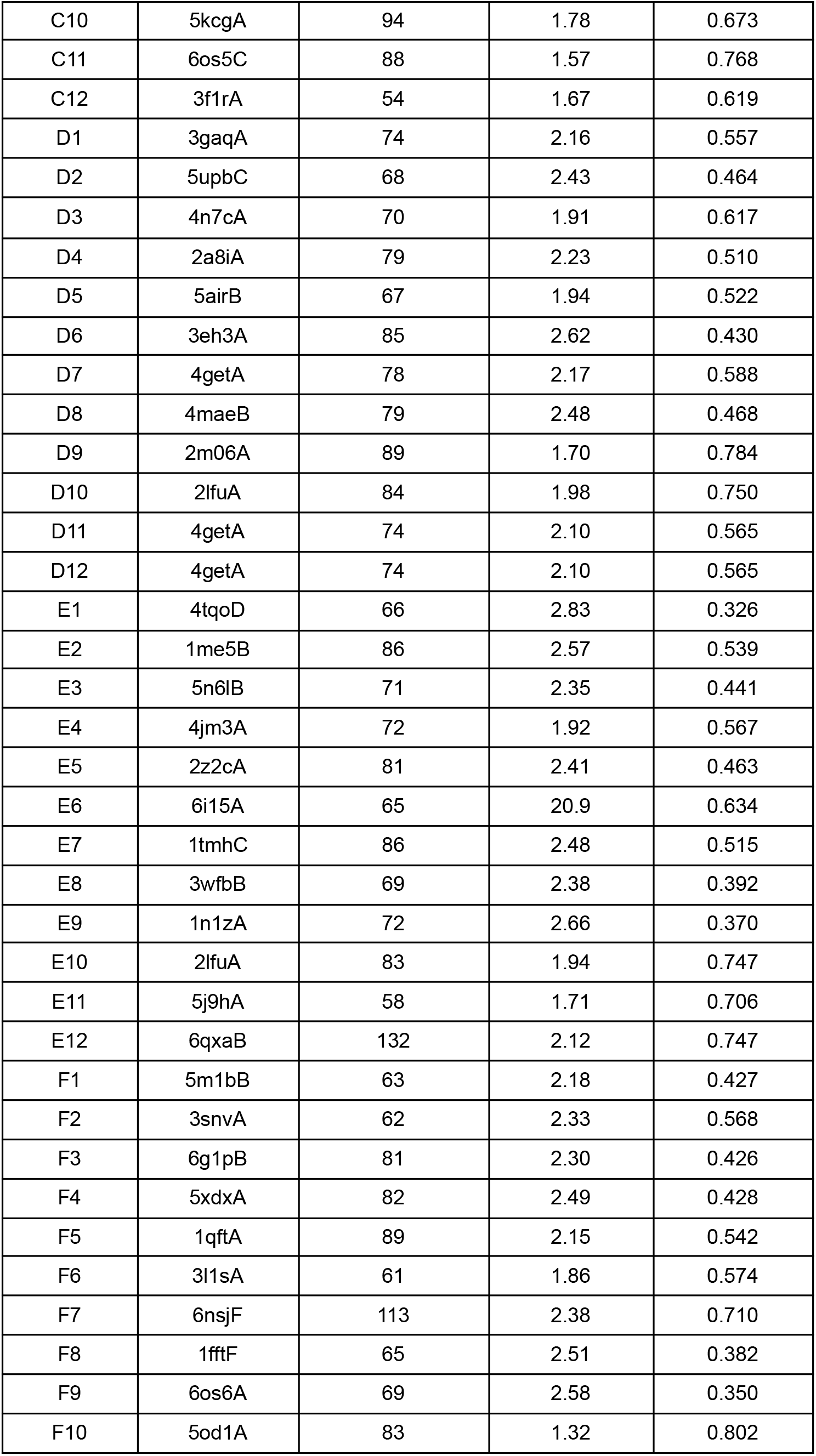
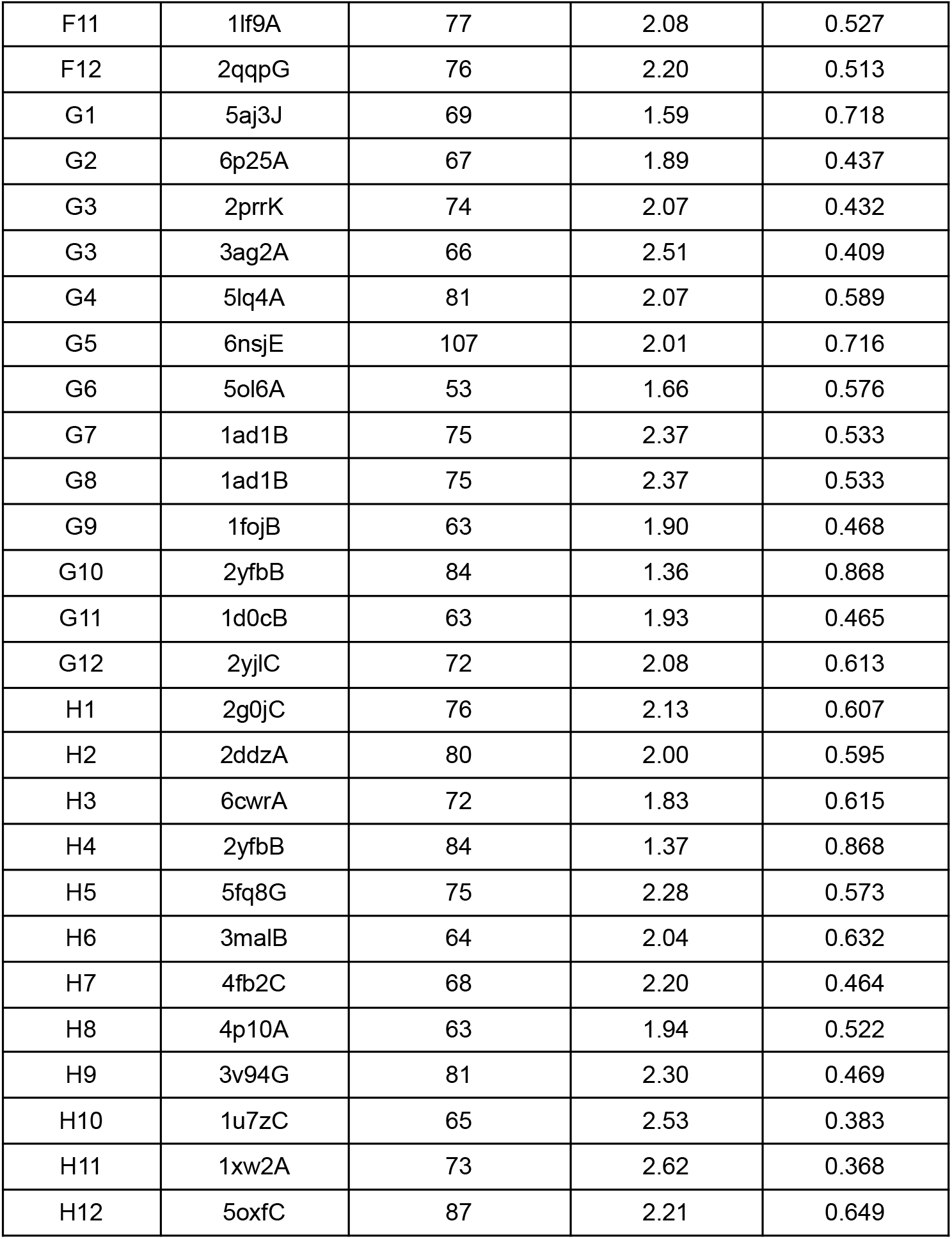
The structural similarity of 96 experimentally-characterized pseudocycles to native proteins in PDB by mTMalign.

**Table S3.**
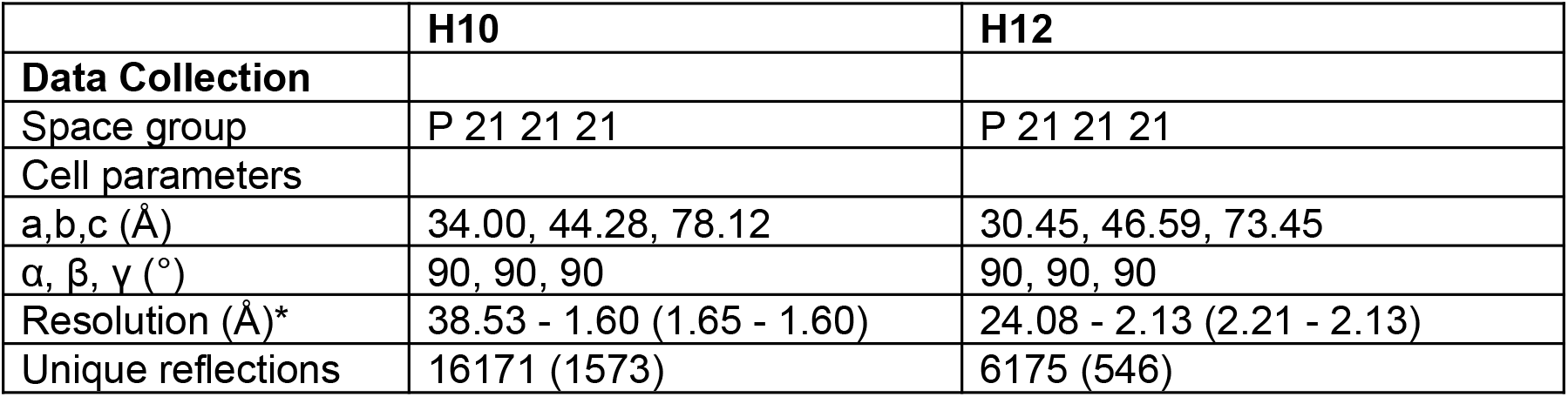
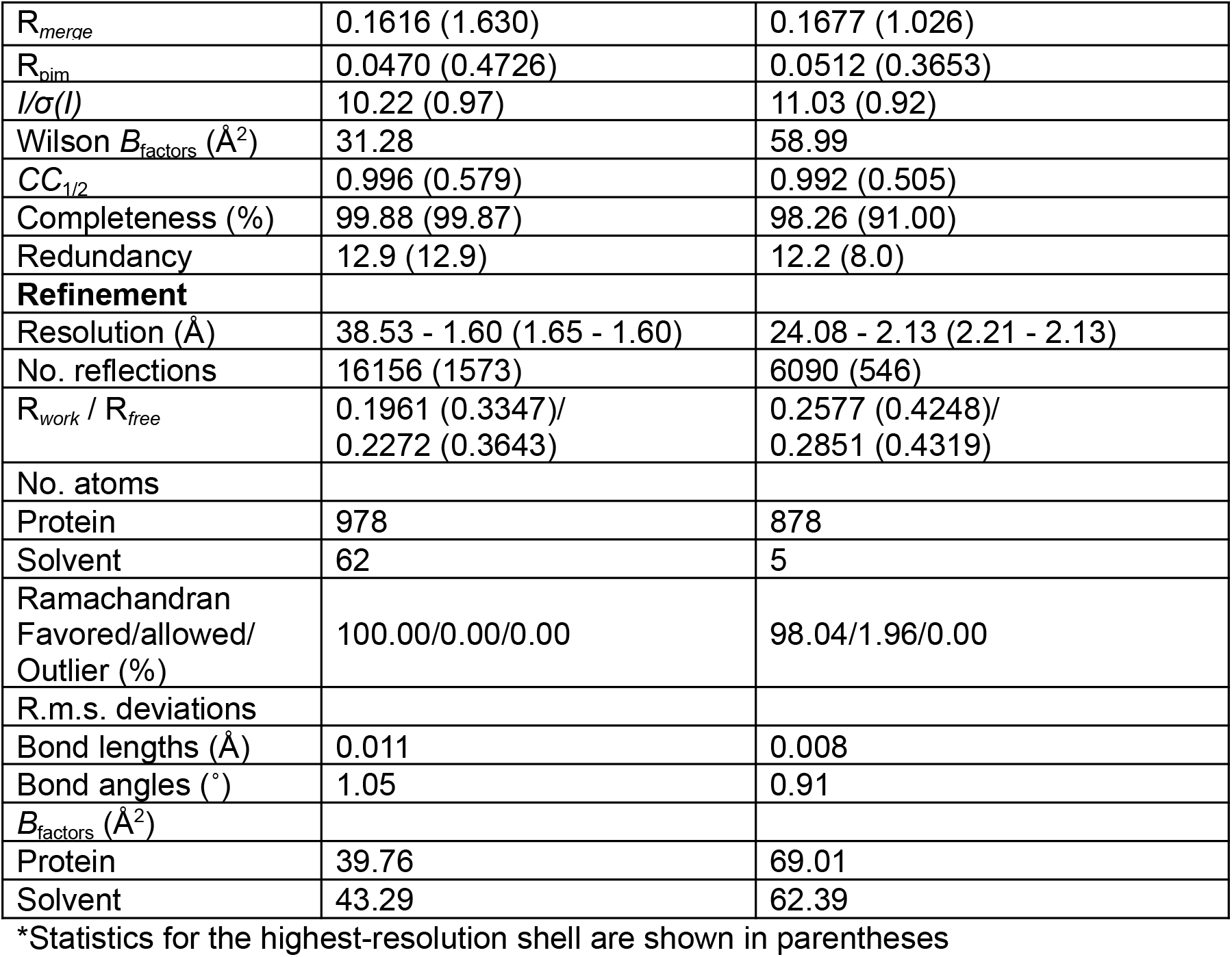
Crystallographic data collection and refinement statistics.

## Notes

### Competing Interest Statement

The authors have declared no competing interest.

